# Therapeutic Eradication of Cancer-associated Fibroblasts Inhibits in vivo progression of Pancreatic Cancer

**DOI:** 10.1101/2025.11.04.686484

**Authors:** Matilda Roos-Mattila, Harri Mustonen, Markus Vähä-Koskela, Sonja Boyd, Sanna Vainionpää, Rosa Hwang, Pauli Puolakkainen, Vadim Le Joncour, Pirjo Laakkonen, Hanna Seppänen

**Affiliations:** Department of Surgery, Helsinki University Hospital, Helsinki; Translational Cancer Medicine Research Program, Faculty of Medicine, University of Helsinki, Helsinki; iCAN Digital Precision Cancer Medicine Flagship, University of Helsinki, Helsinki; Department of Pathology, Helsinki University Hospital, Helsinki; Department Of Surgical Oncology, Division Of Surgery, The University Of Texas MD Anderson Cancer Center, Houston, TX, USA; Neuroscience Center, Helsinki Institute of Life Science (HiLIFE), University of Helsinki, Helsinki, Finland; Laboratory Animal Center, Helsinki Institute of Life Science (HiLIFE), University of Helsinki, Helsinki, Finland

**Author notes:** **Correspondence:** Pirjo Laakkonen, Haartmaninkatu 8, 00290 Helsinki. Equal contribution (Vadim Le Joncour, Pirjo Laakkonen, Hanna Seppänen).

**Keywords:** Pancreatic ductal adenocarcinoma, Cancer-Associated Fibroblast, cationic amphiphilic drug, lysosomal membrane permeabilization, Clemastine, Cancer Medicine Research, Single-cell transcriptomics, Metastasis

## Abstract

Pancreatic ductal adenocarcinoma (PDAC) remains a lethal disease with an unmet medical need. therapeutic elimination of cancer associated fibroblasts (CAFs), key drivers of tumor aggressiveness, has been the focus of recent studies. However, the inherent heterogeneity and plasticity of CAFs have hampered the development of CAF-targeted therapies. Clemastine, an FDA-approved cationic amphiphilic drug, is known to elicit cytotoxic lysosomal membrane permeabilization in solid tumors. We evaluated its efficacy in patient derived PDAC organoids and patient avatars. *In vitro,* half of the organoids responded to clemastine, although sensitivity did not predict in vivo outcomes. *In vivo*, clemastine treatment led to CAF depletion, halted cancer progression and delayed disease progression**. To** assess potential synergy with standard-of-care therapy, clemastine was combined with gemcitabine. The combination enabled dual targeting—clemastine effectively eliminated all CAF subtypes, while gemcitabine eradicated cancer cells - leading to inhibition of metastatic dissemination. These findings support clemastine as a promising companion therapy in PDAC, targeting the tumor-supportive microenvironment.

## INTRODUCTION

Pancreatic ductal adenocarcinoma (PDAC) is notorious for its aggressiveness and very poor prognosis.^1,2^ Several subtypes of cancer-associated fibroblasts (CAFs) form the major component of the especially profound stroma of PDACs. The fibrotic stroma produced by both tumor cells and CAFs renders the tumor about three times stiffer than the healthy pancreatic tissue.^3,4^ Increased tissue stiffness combined with poor vascularization of pancreatic tumors increase hypoxia-driven epithelial-to-mesenchymal transition^5^, limit immune cell infiltration and impair intratumoral drug delivery.^6^

As CAFs and the tumor microenvironment play a pivotal role in PDAC progression, targeting CAFs has become a central focus of recent research. However, due to their heterogenous origin and diverse functions, development of clinical strategies to target CAFs has proven challenging.^7,8^ Several CAF subtypes have been identified based on the expression of distinct marker proteins. One subtype, known as cytokine secreting/inflammatory CAFs (iCAFs), is characterized by the upregulation of the fibroblast activation protein (FAP) and interleukin 6 (IL-6) along with downregulation of the alpha smooth muscle actin (aSMA). iCAFs contribute to the immunosuppressive tumor microenvironment and induce cachexia in PDAC patients.^8^ Another subtype, myofibroblastic CAFs (myCAF, FAP+, aSMA^high^, IL-6^low^), are in close proximity to the neoplastic cells, forming a periglandular ring surrounding cancer cell clusters. Physical contact with tumor cells dictates myCAF adaptation, as *in vitro* studies revealed that stellata cells that do not interact with PDAC cells in co-culture exhibited predominantly iCAF characteristics.^8^

In addition, antigen presenting CAFs (apCAFs) form a smaller subtype in PDACs with anti-inflammatory, tumor supporting features.^7^ In contrast, defensive/tumor suppressive CAFs show tumor-suppressing functions and features.^9^

In response to the nutrient-deprived conditions of the tumor microenvironment, PDAC cells reprogram their metabolic machinery by activating lysosome-dependent pathways.^10^ This metabolic adaptation is closely linked to increased malignancy and reflected in alterations in lysosomal morphology ^11^ and increased cathepsin activity.^12^ These “neoplastic” lysosomes provide several advantages to tumor cells including improved neovascularization and promotion of invasive growth.^13,14^ On the other hand, they also render tumor cells vulnerable to therapy. For instance, increased cathepsin activity is detrimental to lysosome-associated membrane proteins (LAMPs), compromising lysosomal integrity and sensitizing cells to lysosomal membrane permeabilization (LMP) induced cell death.^12^

Thus, lysosomes have emerged as promising therapeutic targets in cancer. Several strategies have been investigated to disrupt lysosomal function such as inhibition of lysosomal hydrolases, heat shock protein 70 and proton pump activity, as well as using chloroquine derivatives.^11^ While many of these approaches have shown efficacy *in vitro*, their clinical translation has been limited due to inconsistent outcomes in PDAC and other cancers.^15-18^ Toxicity and off-target effects of these compounds have been the major complications.^19^ A recently identified alternative for therapeutic lysosomal targeting involves the use of cationic amphiphilic drugs (CADs), compounds that include a wide range of FDA-approved drugs such as antidepressants, neuroleptics, and antihistamines.^20^ CADs possess a dual chemical structure: their cationic charge facilitates diffusion through biological membranes at neutral pH while their amphiphilic moiety becomes protonated in lower pH, leading to entrapment within the lysosomal lumen. CADs are also potent inhibitors of lipid catabolic enzymes, disrupting the normal turnover of lysosomal membrane lipids, and thereby further promoting LMP.^21^ Among CADs, well-tolerated antihistamines have shown cancer cell killing activity in pre-clinical studies and significantly reduced cancer incidence and mortality in epidemiological analyses.^22^

CADs have not been studied in PDAC treatment. Our previous work identified clemastine, an antihistamine CAD, as an effective inhibitor of glioblastoma invasion in vivo.^23^ Considering the shared characteristics of aggressive dissemination between glioblastoma and PDAC, we assessed PDAC sensitivity to clemastine treatment. To this end, we developed research models including patient-derived organoids (POs) and *in vivo* orthotopic murine models, i.e. patient avatars to evaluate the preclinical efficacy of clemastine, gemcitabine, and their combination. Our results show that clemastine effectively eradicated most subtypes of CAFs within the patient avatar tumors, preventing tumor growth and metastatic progression. Using high-throughput cell and tumor secretome profiling as well as single-cell transcriptomics (scRNAseq), we characterized CAF populations at both cellular and molecular levels, uncovering previously unrecognized vulnerabilities unlocked by clemastine.

## RESULTS

### Sensitivity of PDAC organoids to clemastine treatment

To study the efficacy of clemastine on PDACs, we established an orthotopic PDAC mouse model. First, we generated tumor organoid cultures (PO34, PO37, PO77, and PO80) from four PDAC patients operated at the Helsinki University Hospital. We also developed organoids from histologically normal pancreatic tissue adjacent to tumor from one patient (PO34A) (**Fig.1**). The most common PDAC driver mutations, such as Kirsten rat sarcoma virus (KRAS) and tumor suppressor protein p53 (TP53), were detected in all the established tumor POs consistent with the mutational profiles of original tumors for PO34 and PO77.^24^ Mutation data was not available from original tumors for the PO37 and PO80. These mutations were absent in the tumor-adjacent organoid PO34A (**Fig. 2A**). Morphologically, the established tumor POs exhibited typical PDAC organoid architecture whereas PO34A displayed features characteristic for healthy pancreatic ductal organoids (**Fig. 2B**).

**Figure 1.**
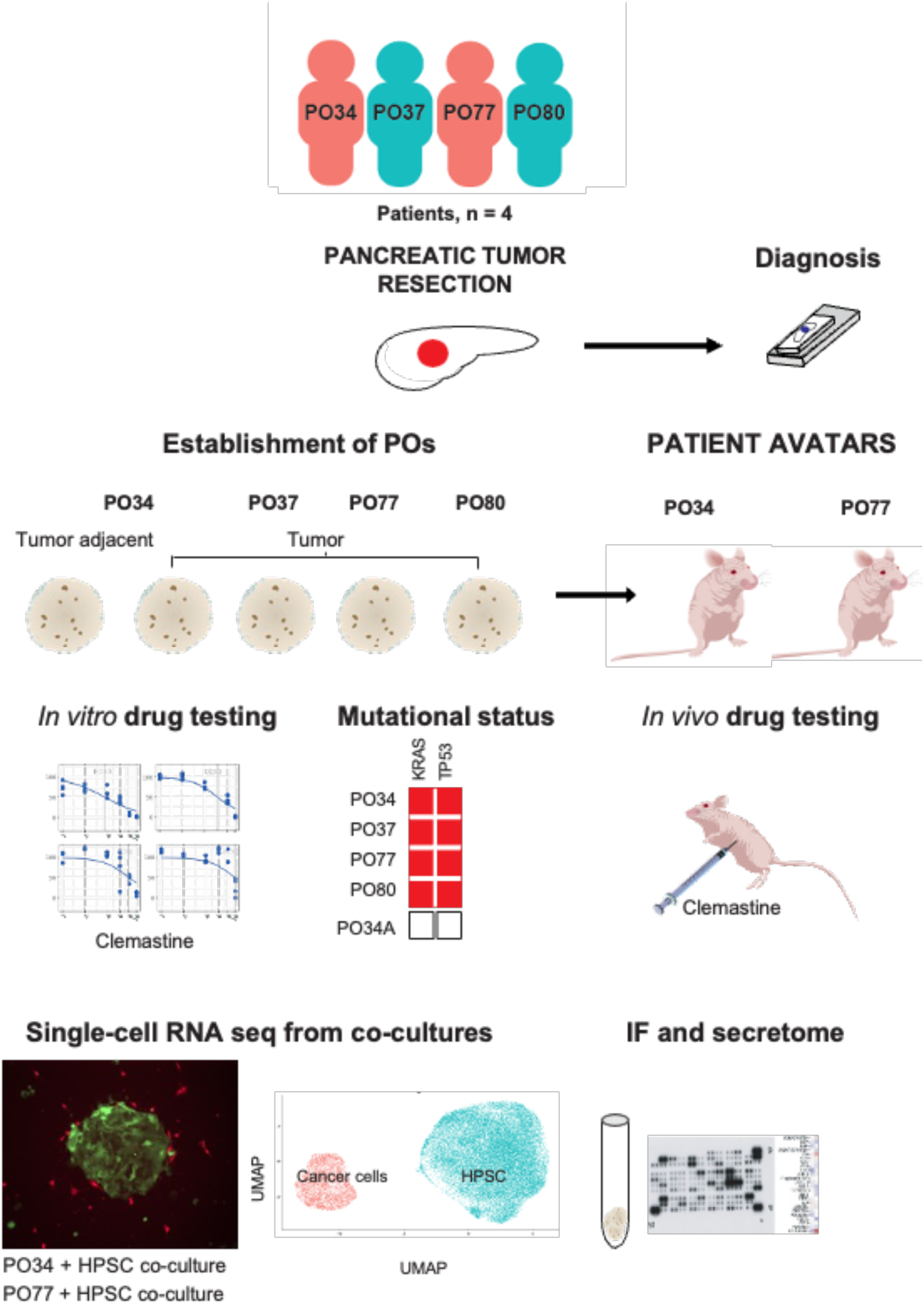
Study protocol. Four patients with PDAC were included in the study. A sample from the resected pancreatic specimen was transferred to the laboratory to grow the tumor POs (PO34, PO37, PO77 and PO80). In addition, one sample of tumor adjacent, histologically normal pancreatic tissue was obtained and is referred to as PO34A. All cell cultures were tested for clemastine sensitivity went through mutational analysis. PO34 (clemastine-sensitive in vitro) and PO77 (clemastine-resistant in vitro) cells were surgically implanted in the pancreas of immunocompromised nude mice, a model referred to as patient avatars. This enabled preclinical evaluation of clemastine as a monotherapy and in combination with the therapy of reference, gemcitabine. To understand the transcriptomic profiles underlying clemastine sensitivity, we performed single-cell RNA sequencing (scRNAseq) on clemastine treated and untreated co-cultures of human pancreatic stellate cells (hPSC) and PO34 (clemastine-sensitive in vitro) or PO77 (clemastine-resistant in vitro). To further examine the effects of clemastine, tumor secretome from PO and patient avatar extracts were analyzed to connect deregulated biological processes with clemastine treatment.

**Figure 2.**
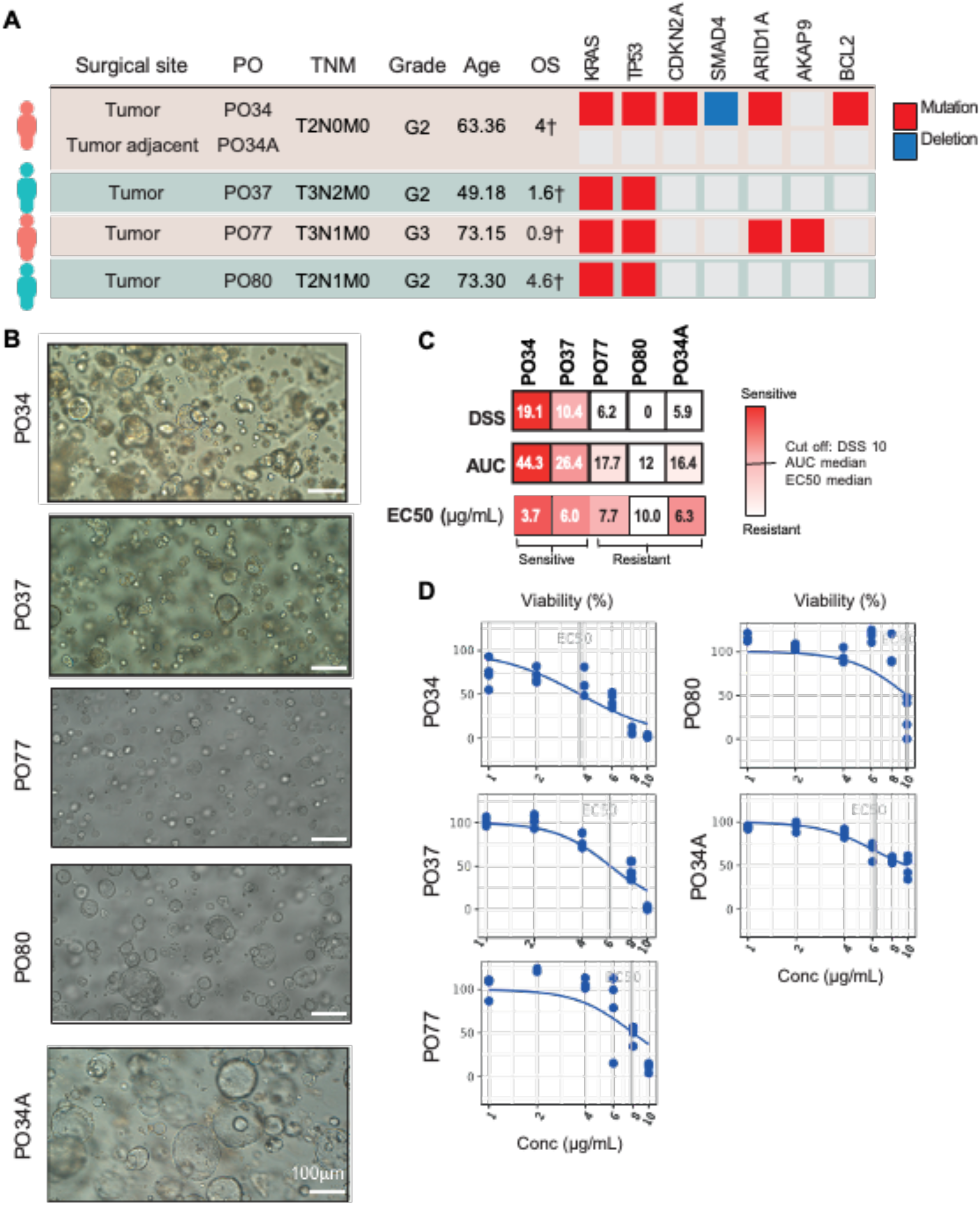
Heterogenous in vitro sensitivity of POs to clemastine. **A.** Patient characteristics: TNM, grade, age (at operation), known somatic mutations and deletions detected in the POs, overall survival (OS). †Deceased. CDKN2A=cyclin-dependent kinase inhibitor 2A, SMAD4=SMAD family member 4, ARID1A=AT-rich interactive domain-containing protein 1A, AKAP9=A-kinase anchor protein 9, BCL2=B-cell lymphoma 2. **B**. Transmitted light micrographs of the established POs. Scale bars 100 µm. **C.** Summary of the DSS (drug sensitivity score), area under the curve (AUC), and half-maximal effective concentration (EC50) values for PO34, PO37, PO77, PO80 and the tumor adjacent PO34A treated with clemastine. PO34 and PO37 showed sensitivity to clemastine treatment, while PO77 and PO80 along with the tumor adjacent PO34A did not respond to clemastine. Sensitivity cutoff: DSS>10. **D**. Clemastine dose response curves for tumor organoid and tumor adjacent POs.

To assess the sensitivity of the POs to clemastine, organoids were treated with increasing drug concentrations (1-10 µg/mL) and the cell viability was assessed on days 1, 3 and 5 post-treatment. Half of the tumor-derived POs (PO34 and PO37) showed significant clemastine sensitivity *in vitro* with drug sensitivity score (DSS) greater than 10,^25^ while PO77, PO80 and the tumor adjacent PO34A were resistant (**Fig. 2C, D**). We also treated the commercial cell lines (MiaPaca-2, HPAF-II, and AsPC-1) with clemastine. Over 50% of the MiaPaca-2 cells were killed at concentrations of 8 and 10 µg/mL of but not at the lower doses (**Supplementary Fig. S1**). HPAF-II cells showed over 50% killing only on day 5 at 10 µg/mL concentration whereas the AsPC-1 cells remained resistant with all tested concentrations (**Supplementary Fig. S1**).

Next, we evaluated potential synergistic effects between clemastine and three approved PDAC treatment regimens (Gemcitabine, Paclitaxel and FOLFIRINOX) as well as one investigational MEK inhibitor (Trametinib). PO34 and PO77 tumor-derived organoids were treated with each drug alone or in combination using fixed concentration of clemastine with increasing concentrations of each drug (**Supplementary Table 1**). Cell viability was assessed at day six post-treatment. No additive or synergistic effects were detected *in vitro* **(Supplementary Fig. S2)**.

### Clemastine inhibits progression of PDAC tumors in vivo

To generate PDAC patient avatars, 5000 PO-derived cells were injected through the splenic vein upstream of the pancreas. In a pilot experiment, we evaluated the *in vivo* efficacy of clemastine using tumors derived from the clemastine-sensitive PO34 organoids. Daily intraperitoneal (IP) injections (3 µL/s) of clemastine or saline as control (n=8 in each group) were started at twenty days (D20) post-implantation (**Supplementary Fig. 3A**). The study was terminated at D60 and clinically relevant tissues from primary and metastatic sites (spleen, liver, and lymph nodes) were harvested for histopathological analysis. Consistent with our previous data in immunocompromised mice, ^23^ the treatment was well-tolerated with no significant body weight differences between treated and control animals (**Supplementary Fig. 3B**).

Histopathological evaluation revealed that the total tumor volume was significantly smaller in the clemastine-treated patient avatars compared to the vehicle group (p = 0.003, **Supplementary Fig. 3C**). All tumors except one resistant outlier responded with substantial size reduction including cases of total tumor remission. Tumors in the vehicle-treated animals exhibited a classical human PDAC histology characterized by multinodular glands in a fibrosis-dense, poorly vascularized stroma (**Supplementary Fig. 3D-F**).^26^

For further histological evaluation, we labeled endothelial cells with antibodies against podocalyxin, a blood vessel marker protein that we previously identified as prognostic biomarker in PDACs.^27,28^ For fibroblast staining, we used antibodies against the glycan NG2, which is highly expressed by the pericytes and a subset of CAFs.^7,29^ Clemastine treatment did not significantly alter the tumor or pancreatic blood vessel densities compared to controls (**Supplementary Fig. 3D-K**). However, vehicle-treated control tumors displayed elevated NG2 expression, while the adjacent pancreatic tissue showed only a few NG2-positive perivascular cells (**Supplementary Fig. 3D-E)**. In contrast, all clemastine-treated tumors, including the resistant outlier, showed a marked reduction in the NG2-positive tumor stroma (**Supplementary Fig. 3G-I**). Interestingly, clemastine selectively affected the NG2-positive tumor stroma as the adjacent healthy pancreas showed no changes in the NG2-positive pericytes or podocalyxin-positive endothelial cells (**Supplementary Fig. 3E**). These results prompted us to further investigate how clemastine affects the cellular composition of the PDAC microenvironment.

### Clemastine-treatment eradicates CAFs in PDAC tumors

CAFs are a key component of the PDAC tumor microenvironment, and they produce most of the extracellular matrix proteins. Moreover, PDAC progression notoriously relies on CAFs.^7^ To further dissect the interplay between clemastine treatment and CAFs, we established two patient avatar models from clemastine-sensitive (PO34) and -resistant (PO77) PDAC organoids. Patient avatars were treated with saline vehicle, clemastine, gemcitabine, a clinically approved drug for PDAC treatment, or a combination of clemastine and gemcitabine starting at D20 post implantation (**Supplementary Fig. 4A-F**). Treatments were well tolerated, and animal body weight did not significantly differ between the groups (**Supplementary Fig. 4B, F**). On D60, treatments were discontinued, and the avatars were sacrificed for histopathological analysis of the pancreas and classical PDAC metastatic sites (spleen, liver, and lymph nodes) (**Supplementary Fig. 4A, E**). Macroscopic pathology during autopsy revealed significant dyspepsia in the gemcitabine and combination treatment groups (**Supplementary Fig. 4C-D**).

Compared to the controls, clemastine treatment led to a significant tumor size reduction in PO77 avatars (p=0.0023), whereas no such effect was detected in the PO34 avatar model (p>0.999) but a trend for tumor size reduction was observed (**Fig. 3A-D**). Tumor sizes varied significantly among the clemastine-treated avatars, a pattern consistently observed across independent experiments and patient-derived models. In 2 out of 7 mice, tumor size reduction was particularly pronounced, while the remaining animals showed variable responses (in accordance with the tumor size heterogeneity detected in the pilot experiment, **Fig. S3C**). Both PO34 and PO77 models responded very well to the gemcitabine monotherapy (PO34 p=0.0516, PO77 p<0.0001) (**Fig. 3B, D**). Due to the marked efficacy of gemcitabine alone, no additional benefit in tumor size reduction was observed in the combination treatment compared to gemcitabine monotherapy (p>0.999) (**Fig. 3B, D**).

**Figure 3.**
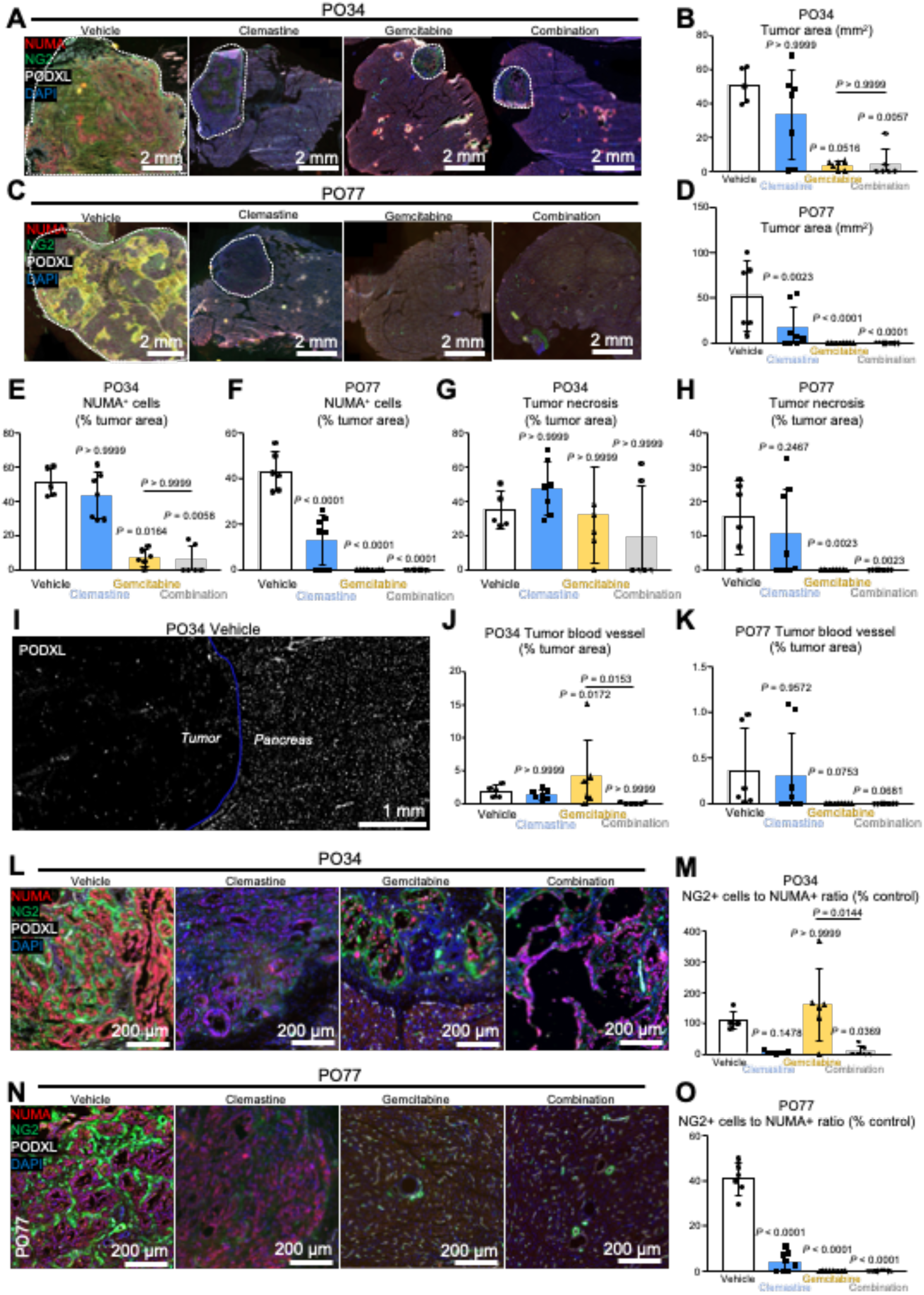
Preclinical assessment of clemastine, gemcitabine and combination regimens in PDAC patient avatars reveals major stromal re-organization associated with clemastine treatment. **A.** Immunofluorescence staining of PO34 PDAC patient avatars in the vehicle, clemastine, gemcitabine, and combination treatment cohorts. Human tumor cells (NUMA, red), cancer-associated fibroblasts (NG2, green), blood vessels (PODXL, white), cell nuclei (DAPI, blue). **B.** PO34 PDAC tumor area in the vehicle (white, N=5), clemastine (blue, 50 mg/kg, N=7), gemcitabine (yellow, 100 mg/kg, N=7), and combination (grey, N=8) cohorts. **C.** Immunofluorescence of PO77 PDAC patient avatars in the vehicle, clemastine, gemcitabine, and combination treatment cohorts. Human tumor cells (NUMA, red), cancer-associated fibroblasts (NG2, green), blood vessels (PODXL, white), cell nuclei (DAPI, blue). **D.** PO77 PDAC area in the vehicle (white, N=6), clemastine (blue, 50 mg/kg, N=9), gemcitabine (yellow, 100 mg/kg, N=9), and combination (grey, N=10) cohorts. **E.** Proportion of PO34 PDAC tumor cells (NUMA+) within the total tumor area in the vehicle (white, N=5), clemastine (blue, 50 mg/kg, N=7), gemcitabine (yellow, 100 mg/kg, N=7), and combination (grey, N=8) cohorts. **F.** Proportion of PO77 PDAC tumor cells (NUMA+) within the total tumor area in the vehicle (white, N=6), clemastine (blue, 50 mg/kg, N=9), gemcitabine (yellow, 100 mg/kg, N=9), and combination (grey, N=10) cohorts. **G.** Quantification of necrotic tissue in the total PO34 PDAC tumor in the vehicle (white, N=5), clemastine (blue, 50 mg/kg, N=7), gemcitabine (yellow, 100 mg/kg, N=7), and combination (grey, N=8) cohorts. **H.** Quantification of necrotic tissue in the total PO77 PDAC tumor in the vehicle (white, N=6), clemastine (blue, 50 mg/kg, N=9), gemcitabine (yellow, 100 mg/kg, N=9), and combination (grey, N=10) cohorts. **I.** Representative micrograph of the distinct vascular endothelium (PODXL, white) in the tumor versus non-neoplastic pancreas (blue line). **J.** Vascular density quantification in the PO34 PDAC (PODXL+) reported to the total tumor area in the vehicle (white, N=5), clemastine (blue, 50 mg/kg, N=7), gemcitabine (yellow, 100 mg/kg, N=7), and combination (grey, N=8) cohorts. **K.** Vascular density quantification in the PO77 PDAC (PODXL+) reported to the total tumor area in the vehicle (white, N=6), clemastine (blue, 50 mg/kg, N=9), gemcitabine (yellow, 100 mg/kg, N=9), and combination (grey, N=10) cohorts. **L.** Visualization of tumor cell (NUMA, red)-associated fibroblasts (NG2, green) in the PO34 PDAC patient avatars of the vehicle, clemastine, gemcitabine, and combination treatment cohorts. Blood vessels (PODXL, white), cell nuclei (DAPI, blue). **M.** Quantification of cancer-associated fibroblasts (NG2+) associated with the PO34 PDAC tumor cells (NUMA+) in the total tumor in the vehicle (white, N=5), clemastine (blue, 50 mg/kg, N=7), gemcitabine (yellow, 100 mg/kg, N=7), and combination (grey, N=8) cohorts. **N.** Visualization of tumor cell (NUMA, red)-associated fibroblasts (NG2, green) in the PO77 PDAC patient avatars in the vehicle, clemastine, gemcitabine, and combination treatment cohorts. Blood vessels (PODXL, white), cell nuclei (DAPI, blue). **O.** Quantification of cancer-associated fibroblasts (NG2+) associated with the PO77 PDAC tumor cells (NUMA+) in the patient avatars in the vehicle (white, N=6), clemastine (blue, 50 mg/kg, N=9), gemcitabine (yellow, 100 mg/kg, N=9), and combination (grey, N=10) cohorts. In the absence of tumors in the gemcitabine and combination groups, CAF quantification could not be performed. *P*-values calculated with 1-way ANOVA and Kruskal-Wallis (B, E, G, J, M) or Holm-Sidak (D, F, H, K, O) post hoc multiple comparison tests.

Quantification of human-specific NUMA-positive tumor cells confirmed the observed tumor volume reduction, showing significant decrease in the PO77 avatars (p<0.0001) and a trend toward reduction in the PO34 avatars following clemastine treatment (**Fig. 3E-F**). Interestingly, both avatar models exhibited prominent necrotic features across all treatment groups (**Fig. 3G-H**). Treatments did not alter the blood vessel distribution or density (**Fig. 3J-K**), classically poor in the PDAC tissue (**Fig. 3I**). In gemcitabine-treated PO34 tumors, occasional inclusions of NUMA-negative, non-neoplastic pancreatic cells were observed showing blood vessel density comparable to the healthy pancreas (**Fig. 3J, Supplementary Fig. 4G-H**). Combination treatment consistently led to complete tumor eradication in PO34 avatars, accompanied by the disappearance of tumor blood vessels (**Fig. 3J-K**) and tumor stroma (**Fig. 3L-O**). In PO77 avatars, gemcitabine and combination treatment groups were excluded from these analyses due to the complete absence of detectable tumor tissue.

We next assessed CAF density and distribution using NG2 as a shared marker for fibroblasts and pericytes.^8^ To better understand how the treatments affected the tumor microenvironment, we normalized the number of NG2^+^ CAFs to the human NUMA^+^ tumor cells, excluding necrotic areas from the analysis. Clemastine treatment markedly reduced the CAF-to-tumor cell ratio in both PO34 and PO77 patient avatars (**Fig. 3L-O**). In contrast, NG2+ CAF populations remained largely unaffected in the gemcitabine-treated PO34 avatars (**Fig. 3L-M**). According to the PO34 data, combination treatment did not compromise the individual efficacy of either clemastine or gemcitabine, as tumors were completely eradicated. This further supports the therapeutic compatibility of the two agents **(Fig. 3N-O).**

PDAC CAFs exist as different subtypes^6-8^. The two main categories include αSMA^high^ peri-glandular myCAFs located close to the tumor cells, and the more distantly located, α-SMA^low^ iCAFs.^8^ Due to the heterogeneity of CAF-populations, we assessed the expression and distribution of several CAF markers including platelet derived growth factor beta (PDGFR-β), α-SMA, NG2 (**Fig. 4A-C, Supplementary Fig. 5A-L**), and FAP (**Supplementary Fig. 5M-N**). Marker expression was correlated to the distance from tumor glands to differentiate myCAFs and iCAFs (**Fig. 4A-C**, **Supplementary Fig. 5**). All labeled CAF subtypes (PDGFR-β p=0.0391, NG2 p<0.001, α-SMA p=0.0488) were negatively affected by the clemastine monotherapy in the PO34 avatars (**Fig. 4D-F**) supporting decreased CAF diversity and abundance. In the PO77 avatars, NG2 and α-SMA expression decreased significantly (p<0.0001 and p=0.018), while a trend towards decreased density was observed in the PDGFR-β expressing cells (p=0.053) (**Fig. 4G-I**).

**Figure 4.**
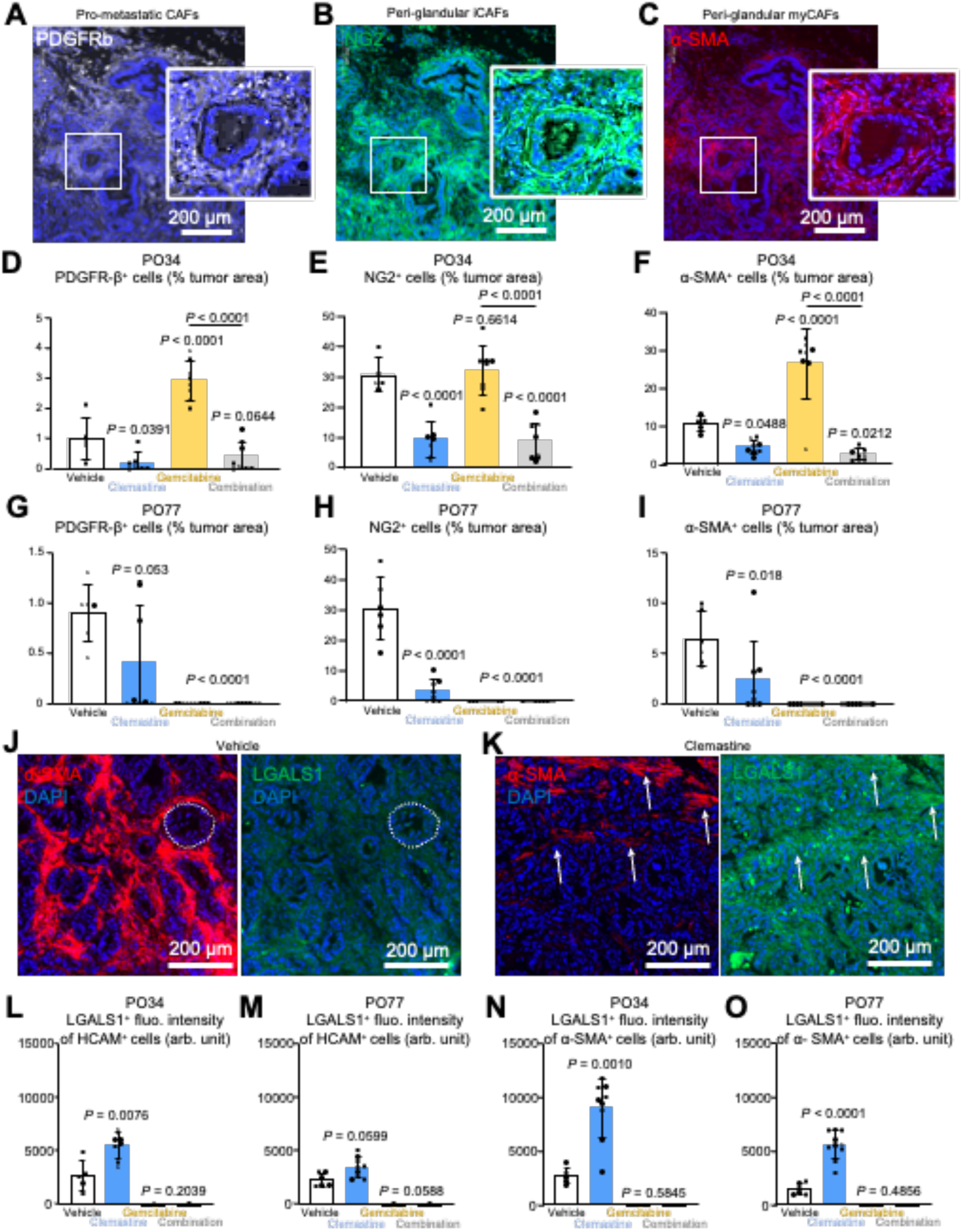
Clemastine treatment depletes PDAC microenvironment from tumor-supporting fibroblasts. **A**. Representative immunofluorescence micrograph of PDGFR-β+ (white) CAFs localized more distant to the tumor glands, typical of iCAF-like phenotype. **B.** Representative immunofluorescence micrograph of NG2+ (green) CAFs surrounding tumor glands, typical of myCAF-like phenotype. **C.** Representative immunofluorescence micrograph of α-SMA+ (red) with α-SMA^high^ CAFs in direct contact with ductal tumor cell glands, typical of myCAF-like phenotype and α-SMA^low^ CAFs more distant to the tumor glands, typical of iCAF phenotype. A-C are co-stained optical fields of a PO77 PDAC control patient avatar, with highlighted magnification of the stromal composition (inserts). Cell nuclei were counterstained with DAPI (blue). **D-F.** Stromal fibroblastic populations in PO34 PDAC patient avatars. Quantification of the PDGFR-β^+^ CAFs in vehicle (white, N=5), clemastine (blue, 50 mg/kg, N=7), gemcitabine (yellow, 100 mg/kg, N=7), and combination (grey, N=8) cohorts. Data expressed as % of the total tumor area (D). Quantification of the NG2^+^ CAFs in vehicle (white, N=5), clemastine (blue, 50 mg/kg, N=7), gemcitabine (yellow, 100 mg/kg, N=7), and combination (grey, N=8) cohorts (E). Data expressed as % of the total tumor area. Quantification of the α-SMA^+^ CAFs in vehicle (white, N=5), clemastine (blue, 50 mg/kg, N=7), gemcitabine (yellow, 100 mg/kg, N=7), and combination (grey, N=8) cohorts (F). Data expressed as % of the total tumor area. **G-I.** Stromal fibroblastic populations in PO77 PDAC patient avatars. Quantification of the PDGFR-β^+^ CAFs in vehicle (white, N=6), clemastine (blue, 50 mg/kg, N=9), gemcitabine (yellow, 100 mg/kg, N=9), and combination (grey, N=10) cohorts. CAFs/tumor area were NQ in gemcitabine and combination treated avatars, as the treatments eradicated tumors completely (G). Data expressed as % of the total tumor area. Quantification of the NG2^+^ iCAFs in vehicle (white, N=6), clemastine (blue, 50 mg/kg, N=9), gemcitabine (yellow, 100 mg/kg, N=9), and combination (grey, N=10) cohorts (H). Data expressed as % of the total tumor area. Quantification of the α-SMA^+^ CAFs in vehicle (white, N=6), clemastine (blue, 50 mg/kg, N=9), gemcitabine (yellow, 100 mg/kg, N=9), and combination (grey, N=10) cohorts (I). Data expressed as % of the total tumor area. **J.** Representative immunofluorescence micrograph of peri-glandular (dotted circle) CAFs expressing α-SMA (red, left panel) and lysosomal permeabilization marker galectin-1 (LGALS1, green, right panel) in a vehicle-treated PO34 PDAC patient avatar. Cytoplasmic LGALS1 fluorescence is distributed evenly in the tumor tissue. **K.** Representative immunofluorescence micrograph of CAFs expressing α-SMA (red, left panel) and lysosomal permeabilization marker galectin-1 (LGALS1, green, right panel) in a clemastine-treated PO34 PDAC patient avatar. Punctate LGALS1 fluorescence is increasingly found in CAFs (arrows). **L.** Quantification of the LGALS1 mean immunofluorescence (arbitrary units) when colocalized with HCAM^+^ ductal tumor cells in PO34 PDAC patient avatars from the vehicle (white, N=5), clemastine (blue, 50 mg/kg, N=7), gemcitabine (yellow, 100 mg/kg, N=7), and combination (grey, N=8) cohorts. **M.** Quantification of the LGALS1 mean immunofluorescence (arbitrary units) when colocalized with HCAM^+^ ductal tumor cells in PO77 PDAC patient avatars from the vehicle (white, N=6), clemastine (blue, 50 mg/kg, N=9), gemcitabine (yellow, 100 mg/kg, N=9), and combination (grey, N=10) cohorts. **N.** Quantification of the LGALS1 mean immunofluorescence (arbitrary units) when colocalized with α-SMA^+^ CAFs in PO34 PDAC patient avatars from the vehicle (white, N=5), clemastine (blue, 50 mg/kg, N=7), gemcitabine (yellow, 100 mg/kg, N=7) and combination (grey, N=8) cohorts. **O.** Quantification of the LGALS1 mean immunofluorescence (arbitrary units) when colocalized with α-SMA^+^ CAFs in PO77 PDAC patient avatars from the vehicle (white, N=6), clemastine (blue, 50 mg/kg, N=9), gemcitabine (yellow, 100 mg/kg, N=9), and combination (grey, N=10) cohorts. NQ in gemcitabine and combination treated avatars, as the treatments eradicated tumors completely. *P*-values calculated with 1-way ANOVA and Kruskal-Wallis (D, E, F, L, M) or Holm-Sidak (G, H, I, M, O) post hoc multiple comparison tests. A, B, C, J, K cell nuclei counterstained with DAPI (blue).

In the gemcitabine-treated tumor remnants, most CAF populations could be found forming “empty sleeves” around the severely disrupted tumor structures consistent with the initial characterization (**Fig. 3L-O**). This indicates that gemcitabine elicited direct tumor cell death and did not exert its antitumoral effects through microenvironmental remodeling. On the contrary, gemcitabine monotherapy led to increase in PDGFR-β^+^ and α-SMA^+^ CAFs per tumor area in the PO34 avatars (<0.0001 for both markers) (**Fig. 4D, F**), but not in NG2^+^ CAFs (p=0.6614) (**Fig. 4E**).The higher CAF-to-tumor cell ratio in gemcitabine-treated PO34 avatars (**Fig. 4D-F**) likely reflects from quantification bias caused by eradication of the neoplastic cells and/or a chemo-induced fibrotic reaction leading to PDGFR-β^+^ and α-SMA^+^ CAF proliferation. Nonetheless, when combined, gemcitabine and clemastine acted synergistically, inducing tumor cell death together with simultaneous decrease in all CAF subtypes (p<0.0001 for all markers, **Fig. 4D-F**).

As both gemcitabine monotherapy and the combination treatment showed very robust anti-tumor effect in the PO77 avatars, no tumor stromal tissue remained for CAF quantification (**Fig. 4G-I)**. However, clemastine retained its efficacy to reduce CAF populations when compared to vehicle treated group (**Fig. 4G-I**) alone or when combined with gemcitabine.

### Clemastine induces lysosomal stress in CAFs

The shift in galectin 1 (LGALS1) localization from diffuse cytoplasmic immunofluorescence to bright punctate lysosomal staining is used to assess lysosomal damage.^30^ In vehicle-treated avatars a faint and diffuse LGALS1 staining was detected (**Fig. 4J**), whereas clemastine treatment increased the punctate staining predominantly in areas containing CAFs (**Fig. 4K**).

We quantified localization of LGALS1 puncta in PDAC cells and CAFs using human homing cell adhesion molecule (hCAM aka CD44) staining to identify tumor cells and α-SMA staining to mark CAFs. This confirmed a significant increase in LGALS1 staining intensity in the PO34 hCAM^+^ tumor cells (p=0.0076) but not in the PO77 tumor cells (p=0.0599) (**Fig. 4L-M**). This suggests that the tumor cells might have maintained the clemastine-sensitive (PO34) and - resistant (PO77) profiles first detected *in vitro*. The clemastine-induced significant increase in the LGALS1 intensity in α-SMA^+^ CAFs was however evident in both PO34 and PO77 (p=0.0010 and p<0.0001) patient avatars and was more pronounced than the increase observed in tumor cells (**Fig. 4N-O**). These results support clemastine-induced lysosomal membrane permeabilization (LMP) in CAFs.

### CAF activation increases their vulnerability to clemastine

Clemastine promoted LMP in CAFs rather than in tumor cells *in vivo*, suggesting selective cell type specific sensitivity. To validate our observations, we applied clemastine to human fibroblasts (WS-1 cell line) and observed a dose-dependent decrease in cell viability (**Fig. 5A**) as well as shift in LGALS1 localization from cytoplasmic to lysosomal (**Fig. 5B-C**) as an indication of lysosomal damage. *In vivo*, CAFs showed marked sensitivity to clemastine, whereas fibroblast/pericyte coverage in the tumor-adjacent pancreas remained unaffected (**Supplementary Fig. 3D-E, J-K**). PDAC CAFs derive from tissue-resident fibroblasts, tumor-infiltrating mesenchymal cells, and pancreatic stellate cells (PSCs).^31,32^ As a preclinical model, we serum-starved PSCs to induce a CAF-like phenotype that activates cancer cell migration.^32^ To study whether this induction would enhance clemastine sensitivity, we treated WS-1 fibroblasts, an immortalised human PSCs line (HPSCs)^33^, and a PDAC-derived CAF cell line (CAF82, isolated in house) with clemastine.

**Figure 5.**
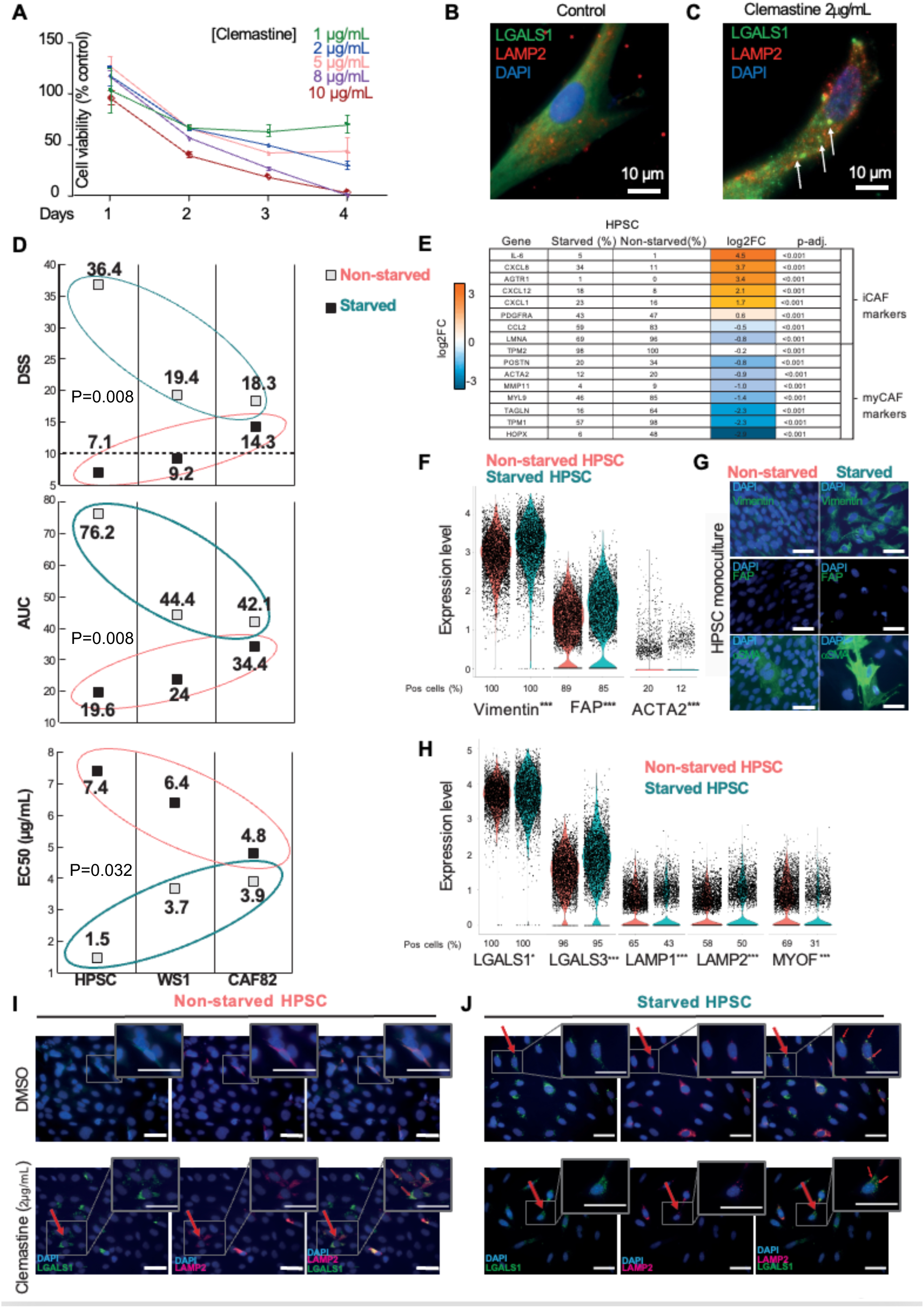
Fibroblast/pancreatic stellate cell sensitivity to clemastine treatment increases with fibroblast activation. **A.** WS-1 human fibroblast viability for indicated clemastine concentrations and timepoints. **B-C.** Representative Immunofluorescence micrographs of control untreated (B, N=10) and clemastine treated (2 µg/mL, 18 h, N=10) (C) WS-1 human fibroblasts. Lysosomal LAMP2 (red) and LGALS1 (green) are visualized. LAMP2^+^/LGALS1^+^ permeable lysosomes are identified by arrows. Cell nuclei counterstained with DAPI (blue). Scalebar: 10 µm. **D.** Clemastine DSS, AUC and EC50 scores in three fibroblast lines (hPSC, WS1, CAF82) in their starved and non-starved states. **E.** Table of putative myCAF/iCAF marker genes and the percentages of cells positive for these in the starved and non-starved HPSC cultures. Average (avg) log2 fold change (FC) for the transcription level differences in activated iCAF-like cells versus parental HPSCs. Adjusted p-values. The change in transcription level (avg log2FC) ranges from blue (downregulated) to orange (upregulated). **F.** Violin plots of scRNAseq gene expression levels of CAF markers vimentin, FAP, and ACTA2 (gene coding for the α-SMA protein) in parental, non-starved (red) and starved, iCAF-like transformed, hPSCs (turquoise). **G.** Representative micrographs of FAP, vimentin, and α-SMA in non-starved (red) and starved (turquoise) HPSCs visualized by IF staining. **H**. Violin plots of scRNAseq gene expression levels of lysosomal markers in non-starved (red) and starved (turquoise) HPSCs. **I-J**. IF staining of LGALS1 (green) and LAMP2 (red) in untreated (DMSO) and clemastine-treated non-starved (I) and starved (J) HPSCs. LAMP2^+^/LGALS1^+^ permeable lysosomes are identified by arrows. Clemastine treatment causes cytoplasmic LGALS1 to relocate to damaged lysosomes. Clemastine treatment also upregulated LAMP2 in HPSCs, but not in iCAF-like cells. LGALS1 relocation is also observed in untreated iCAF-like cells. *p<0.05, **p<0.001, ***p<0.0001.

Induction of CAF-like phenotype by serum starvation consistently enhanced clemastine sensitivity as judged by an increase in both the drug sensitivity score (DSS) and area under the curve (AUC) and accompanied by a corresponding decrease in the half maximal effective concentration (EC50) (**Fig. 5D).** We run multiple replicates in HPSC and found that the starvation-induced increase in all these parameters was statistically significant (p= 0.008, 0.008 and 0.032, respectively). Interestingly, the PDAC-derived CAF cell line (CAF82) exhibited the highest clemastine sensitivity in its non-starved state (DSS 14.3), and the sensitivity was only moderately increased by starvation (DSS 18.3). This confirmed the CAF-like phenotype of CAF82 cells, which was maintained *in vitro* following their isolation from patient tissue.

To study the phenotypic state changes induced by starvation, we performed a scRNAseq analysis from control and starved HPSCs. When using global expression patterns of myCAF and iCAF markers,^8,34^ we observed a downregulation of the myCAF-associated genes upon starvation along enrichment of the iCAF signature (**Fig. 5E**). This suggests that serum starvation of HPSCs preferentially induces an iCAF-like phenotype. Violin plots of CAF marker gene expression - vimentin, FAP, and α-SMA (encoded by *ACTA2* gene) - show upregulation of vimentin and FAP expression in starved cells compared to the non-starved controls (**Fig. 5F**). Based on the scRNAseq data, 20% of control HPSCs expressed α-SMA, while only 12% of the starved cells showed *ACTA2* expression (log2FC -0.9, p=0.0002). Immunofluorescence staining revealed marked upregulation of vimentin protein in starved cells along with increased α-SMA protein expression in a restricted subset of cells. In contrast, FAP protein levels remained unchanged between starved and control cells (**Fig. 5G**).

We further hypothesized that serum starvation-induced iCAF-like phenotype of HPSCs would be accompanied by neoplastic transformation of lysosomes. A significant upregulation of genes involved in lysosomal stress response (*LGALS3*) or disturbances in lysosomal membrane homeostasis (*LAMP1*) was consistently detected following HPSC serum starvation (**Fig. 5H**). No change was observed in the expression *LGALS1*, which is expressed in all cell types, or *LAMP2* levels upon starvation (**Fig. 5H**). Myoferlin (*MYOF*), a membrane-protective lysosomal protein^35^, was significantly downregulated following HPSC serum starvation (p<0.0001) (**Fig. 5H**). Accordingly, punctate staining of the LGALS1 was detected in the lysosomes of serum-starved HPSCs but not in the non-starved controls (**Fig. 5I, J**), indicating acquired lysosomal stress and fragility. The extent of lysosomal stress/damage in starved HPSCs was comparable to that induced by clemastine, which also increased LGALS1 lysosomal puncta formation and upregulated LAMP2 independently of starvation (**Fig. 5I, J**). The serum-starved HPSCs, however, did not further upregulate LAMP2 following clemastine treatment (**Fig. 5J**). Both the lack of additional LAMP2 upregulation, and the reduced *MYOF* expression in the starved HPSCs may explain their elevated susceptibility to clemastine-induced cell death.

### Clemastine reshapes PDAC microenvironment through secretome modulation

To better understand how clemastine cytotoxicity on CAFs could affect PDAC growth, conditioned media from clemastine-treated (2 µg/mL, 24 h) and untreated tumor-adjacent CAF82 cells were analyzed using human cytokine arrays (**Supplementary Fig. 6A-E**). When cytokines were grouped according to their cellular functions, we observed an increased release of over fifteen interleukins following clemastine exposure, including the iCAF-specific IL-6 (**Fig. 6A**). Clemastine treatment also appeared to enhance the immunoreactivity and chemotaxis of CAF82 cells, further supporting the phenotypic similarities between serum starvation-induced iCAF-like transformation of HPSCs and clemastine exposure (**Fig. 6A**). Altogether, the data suggests that short exposure to clemastine induces an inflammatory-like stress response in CAF82 cells.

**Figure 6.**
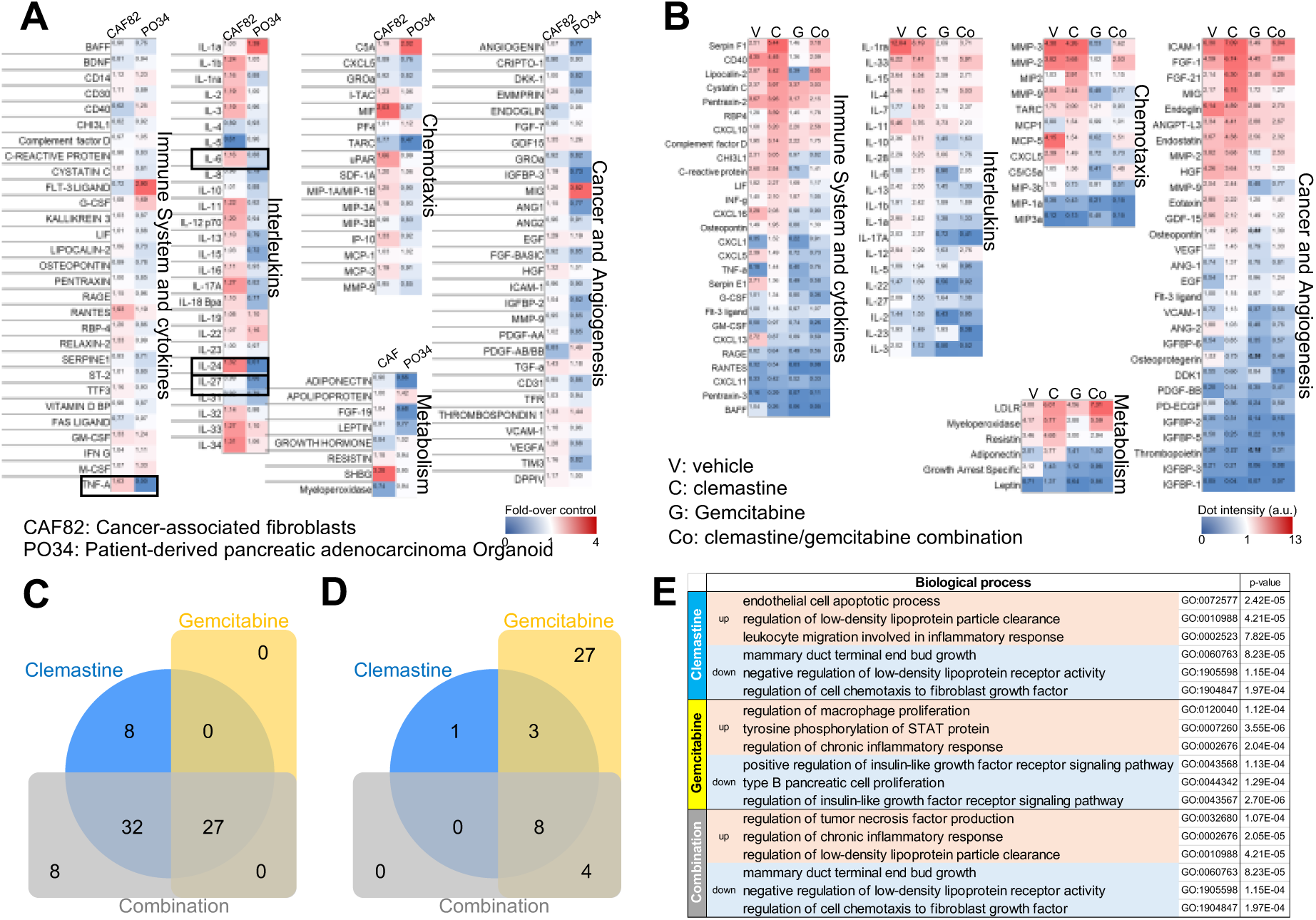
Tumor and CAFs secretomes are modulated by clemastine, gemcitabine or combination regimens. **A.** In vitro human secretome heatmap of conditioned media from human CAF82 (N=2) and PO34 (N=2) organoids treated with clemastine (2 µg/mL, 24 h) for identified groups of cytokines e.g., immune system and cytokines, interleukins, chemotaxis, metabolism, cancer progression, and tumor angiogenesis. Data are expressed as mean of fold-over control untreated cells. **B.** In vivo murine secretome heatmap of PO77 PDAC patient avatar extracts from the vehicle (V), clemastine (C), gemcitabine (G), and combination (Co) treatment groups. Data are expressed as mean protein dot intensity value (arbitrary unit) normalized to assay reference dots. **C.** Venn diagram of downregulated cytokines (ratio cut-off: 0.8) in PO77 PDAC patient avatars in clemastine (blue), gemcitabine (yellow) and combination (grey) groups normalized to the vehicle treatment group. **D.** Venn diagram of upregulated cytokines (ratio cut-off: 1.2) in PO77 PDAC patient avatars in the clemastine (blue), gemcitabine (yellow) and combination (grey) groups normalized to the vehicle treatment group. **E.** Predicted main up- (light red) and down regulated (light blue) biological processes from the secretome of PO77 PDAC patient avatars of the clemastine, gemcitabine and combination treatment groups normalized to the vehicle group.

Under similar conditions, PO34 organoids (**Supplementary Fig. 6A-E**) exhibited a very different response. Globally, clemastine exposure decreased production of almost all analyzed cytokines (**Fig. 6A**). Notably, clemastine exposure led to a strong decrease in secreted tumor necrosis factor alpha (TNF-α), IL-24, and -27. Increased secretion of monokine induced by interferon gamma (MIG), complement component 5A (C5A), IL-1a, and FMS-like tyrosine kinase 3 (FLT-3) may indicate an early paracrine stress response that promotes cancer cell stemness and plasticity (**Fig. 6A**). This stress response was further supported by decreased metabolic, pro-tumoral and angiogenic signals (**Fig. 6A**).

Patient avatar microenvironment composition was screened by using mouse-compatible arrays. To better understand the long-term (40+ days) effect of the treatments, total tumor extracts from the different treatment groups were analyzed (**Supplementary Fig. 6F-J,** **Fig. 6B**). Consistent with the *in vitro* findings, clemastine-treated tumors exhibited increased inflammatory and stress response signals (**Fig. 6B-C**), accompanied by potential upregulation of lipid metabolism, as judged by overrepresentation of low-density lipid receptor (LDLR), myeloperoxidase, and adiponectin (**Fig. 6B-D**). In accordance with the histopathological analysis of the gemcitabine and combination-treated patient avatars, PDAC cell destruction was associated with the marked suppression of cancer-associated signals (**Fig. 6B-D**). Gene ontology (GO) analysis of the biological processes associated with the PDAC microenvironment revealed different stress responses induced by clemastine and gemcitabine. Clemastine primarily disrupted lipoprotein homeostasis, consistent with the classical profile of CADs, whereas gemcitabine affected pancreatic functions (**Fig. 6E**). When administered in combination, gemcitabine and clemastine appear to act synergistically inducing concurrent cytotoxic, proinflammatory, and lipid metabolism disturbances (**Fig. 6E**).

### Molecular landscapes of HPSCs and PDAC cells in response to serum-starvation, co-culture, and clemastine treatment

To generate a comprehensive map of the gene alterations induced by HPSC serum starvation and clemastine treatment as well as to illuminate the intricate relationships between different types of fibroblasts and PDAC cells, we created co-cultures of clemastine sensitive (PO34) and resistant (PO77) PDAC organoids with control HPSCs or serum-starved HPSCs, (referred to as activated, iCAF-like HPSCs) (**Fig. 7A**). Co-cultures were either grown in normal cell culture medium (with or without serum) or in medium supplemented with clemastine (2 µg/mL) for three days. Cell hashing of co-cultures was performed using antibody tagging prior submitting to multiplexed scRNAseq. Individual transcriptomes of around 50 000 cells were characterized and annotated based on their HPSC or ductal tumor cell features using *Seurat* cell clustering and UMAP data projection (**Fig. 7B**). We identified robust canonical PDAC cell markers including *KRT19, SOX9, EPCAM, CD44, CD9, and BPIFBI* (**Fig. 7C**). Expression and occurrence of these genes remained unaffected by the treatment or culture conditions and were undetectable in all HPSC subtypes (**Fig. 7D-E**). Next, we analyzed expression of markers in HPSCs expanded in two-dimensional cultures and assessed how their transcriptomic identity is maintained upon serum starvation, clemastine exposure, or in 3D co-culture with PDAC cells (**Fig. 7F-G**). We identified five strong marker genes in control HPSCs, *COL1A1*, *TAGLN*, *MYL9*, *CXCL1* and *-8* (**Fig. 7F**). Expression of these genes was downregulated following HPSC activation by serum starvation (**Fig. 7F**), consistent with the transcriptional profile of activated 3D HPSCs. The consistent transcriptomic alignment observed across all activated serum-starved HPSC types indicates a robust differentiation process, in line with the pronounced shift towards iCAF-like phenotype shown in **Figures 5 and 6**. Using gene ontology analysis (**Table 1**) and curated pathway enrichment data (**Fig. 7G**), we confirmed that both HPSC activation by serum starvation and clemastine exposure triggered a dramatic cellular stress response and upregulated lipid metabolism activity (**Table 1**). Among the differentially expressed genes (DEGs) with the highest specificity to iCAF-like phenotype following clemastine exposure, *NPC1* (Intracellular Cholesterol Transporter 1), *FABP3 (*Fatty Acid Binding Protein 3), and *HMGCS1* (3-Hydroxy-3-Methylglutaryl-CoA Synthase 1), have previously been associated with lysosomal dysfunction (**Supplementary Fig. 7A-C**).^23,36^

**Figure 7.**
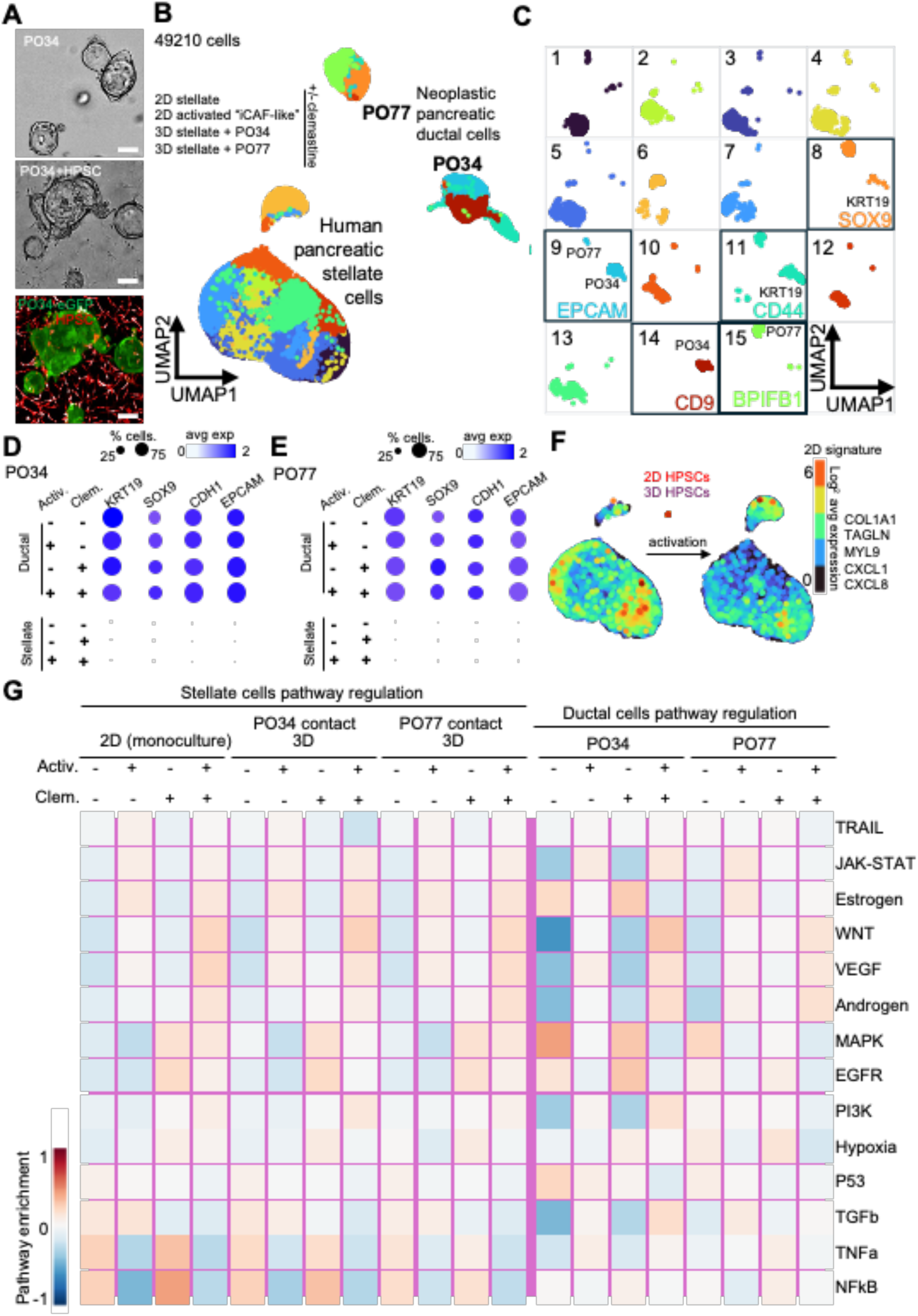
Impact of stellate cell activation, stellate-ductal tumor cell interaction and clemastine treatment on the single cell transcriptome profiles and intracellular network regulation. **A**. Representative brightfield micrograph of monocultured PO34 (upper), co-cultured with HPSC (middle) and fluorescent micrograph of PO34-GFP (green) co-cultured with HPSC-mcherry (red, lower panel). Scale bars 100µm. **B**. Integrated Uniform Manifold Approximation and Projection for Dimension Reduction (UMAP) plot of 49210 barcoded cells including parental human pancreatic stellate cells (HPSCs) and serum-starved activated HPSCs 2D cultures, PO34 and PO77 co-cultured with either HPSCs or serum-starved activated HPSCs, in the absence or presence of clemastine (4 µg/mL, 24h). **C.** Individual cell clusters computed in Seurat. Squared clusters contain ductal neoplastic cells, including cluster 8 upregulating keratin 19 (KRT19)+SOX9, cluster 9 upregulating the epithelial cell adhesion molecule (EPCAM), cluster 11 upregulating KRT19+CD44, cluster 14 upregulating CD9, and cluster 15 upregulating BPI fold-containing family B member 1 (BPIFB1). **D.** Dot plot for PO34 pancreatic ductal tumor cell hallmark genes (as defined in C). Expression of KRT19, SOX9, CDH1 and EPCAM is consistent in ductal but undetectable in stellate cells, regardless of stellate cell activation by serum-starvation (Activ.) or clemastine treatment (Clem.). **E.** Dot plot for PO77 pancreatic ductal tumor cell hallmark genes (as defined in C). Expression of KRT19, SOX9, CDH1 and EPCAM is consistent in ductal but undetectable in stellate cells, regardless of stellate cell activation by serum starvation (Activ.) or clemastine treatment (Clem.). **F**. Gene expression signature distribution of parental (left) and activated by serum-starvation (right) HPSCs (expressed as log2 average gene expression from black to red). Gene signature including upregulated genes in 2D parental HPSCs exhibiting downregulation following HPSC activation by serum-starvation or co-culture in 3D matrices with neoplastic ductal cells, e.g., collagen 1A1 (COL1A1), transgelin (TGLN), myosin regulatory light polypeptide 9 (MYL9), CXCL1 and 8. **G**. Heatmap of main molecular pathway regulation in stellate and ductal tumor cells in response to HPSC activation by serum-stravation (Activ.) and/or clemastine treatment (Clem.). HPSCs were grown in 2D monocultures or in Matrigel co-cultures with either PO34 or PO77 ductal cells. Ductal cells were grown in Matrigel in presence of either parental or serum-starvation activated HPSCs. Pathways were identified in CollecTRI and curated from DoRothEA, TRUST and SIGNOR in deCoupleR running in Bioconductor.

In control HPSCs, angiogenic cell migration and cell adhesion processes were downregulated following clemastine exposure (**Fig. 7G**), in accordance with the secretome data. In-depth pathway enrichment analysis revealed enhanced cholesterol and lipid processing upon HPSC activation, as evidenced by the upregulation of estrogen and androgen signaling pathways (**Fig. 7G**). While clemastine exposure increased hypoxia pathways, it was not followed by increased *VEGF* and *PI3K* activation classically triggered by oxygen deprivation (**Fig. 7G**). Together with TRAIL downregulation and upregulation of *TNFa* and *NFkB* in response to clemastine exposure, these findings suggest a pronounced cellular stress and death response induced by the treatment (**Fig. 7G**). By integrating PDAC transcriptomes into the pathway analysis, we discovered that both PO34 and PO77 exhibited a nearly identical response to serum-starved HPSC, clemastine treatment, and their combination (**Fig. 7G**). For instance, activation of MAPK and EGFR pathways was enhanced when PDAC cells were in direct contact with activated serum-starved HPSCs or exposed to clemastine. Interestingly, this upregulation is buffered when serum-starved iCAF-like, activated HPSCs are added to the PDAC cultures, even in the presence of clemastine (**Fig. 7G**). Pathways responsible for the initiation and progression of PDACs, e.g., WNT and TGFß, are consistently upregulated in presence of the iCAF-like, activated HPSCs. These pathways are partially downregulated upon clemastine exposure (**Fig. 7G**). In these experiments, clemastine exposure was rather short, while data from the secretome and patient avatars suggest that longer exposure to clemastine can eradicate tumor-supporting fibroblasts and overcome PDAC chemoresistance.

### CAF subtypes in response to serum-starvation induced activation and clemastine treatment

To further validate clemastine’s ability to eliminate tumor shielding fibroblasts, we profiled CAF subpopulations and assessed their relative abundance upon serum starvation and/or clemastine exposure. To this end, we plotted the scRNAseq data (**Fig. 7B**) as T-SNE projections to better separate cellular subtypes. These projections revealed a robust expression of PDAC cell signatures both in PO34 and PO77, regardless of the HPSC activation by serum starvation or clemastine exposure (**Fig. 8A**). To focus our analysis on CAF subtypes, we first used the published HPSC iCAF/myCAF/apCAF, and defCAF gene signatures ^7-9,34^ to mark HPSCs in control, activated, and clemastine-treated conditions (**Fig. 8B**). We detected an enrichment of iCAF signatures following HPSC activation (**Fig. 8B**), particularly within activated HPSC populations that showed upregulated IL-6 expression (**Fig. 8C**). When we quantitated the average gene signature enrichment and cell abundance, we observed that HPSC activation increased iCAF cell scores. Interestingly, iCAF signature score and abundance of iCAF-like cells markedly reduced following clemastine treatment, especially when HPSCs had been primed towards iCAF phenotype prior to clemastine exposure (**Fig. 8D**). Further analysis revealed that clemastine exposure led to reduction in both signature scores (**Fig. 8E,G**) and cell abundances (**Fig. 8F,G**) of myCAFs (**Fig. 8E-F**), and to a lesser extent apCAFs (**Fig. 8G-H**). These effects were even more pronounced following HPSC activation. However, gene signature score and cell abundance of defCAFs remained largely unchanged in response to either HPSC activation or clemastine treatment (**Fig. 8I-J**).

**Figure 8.**
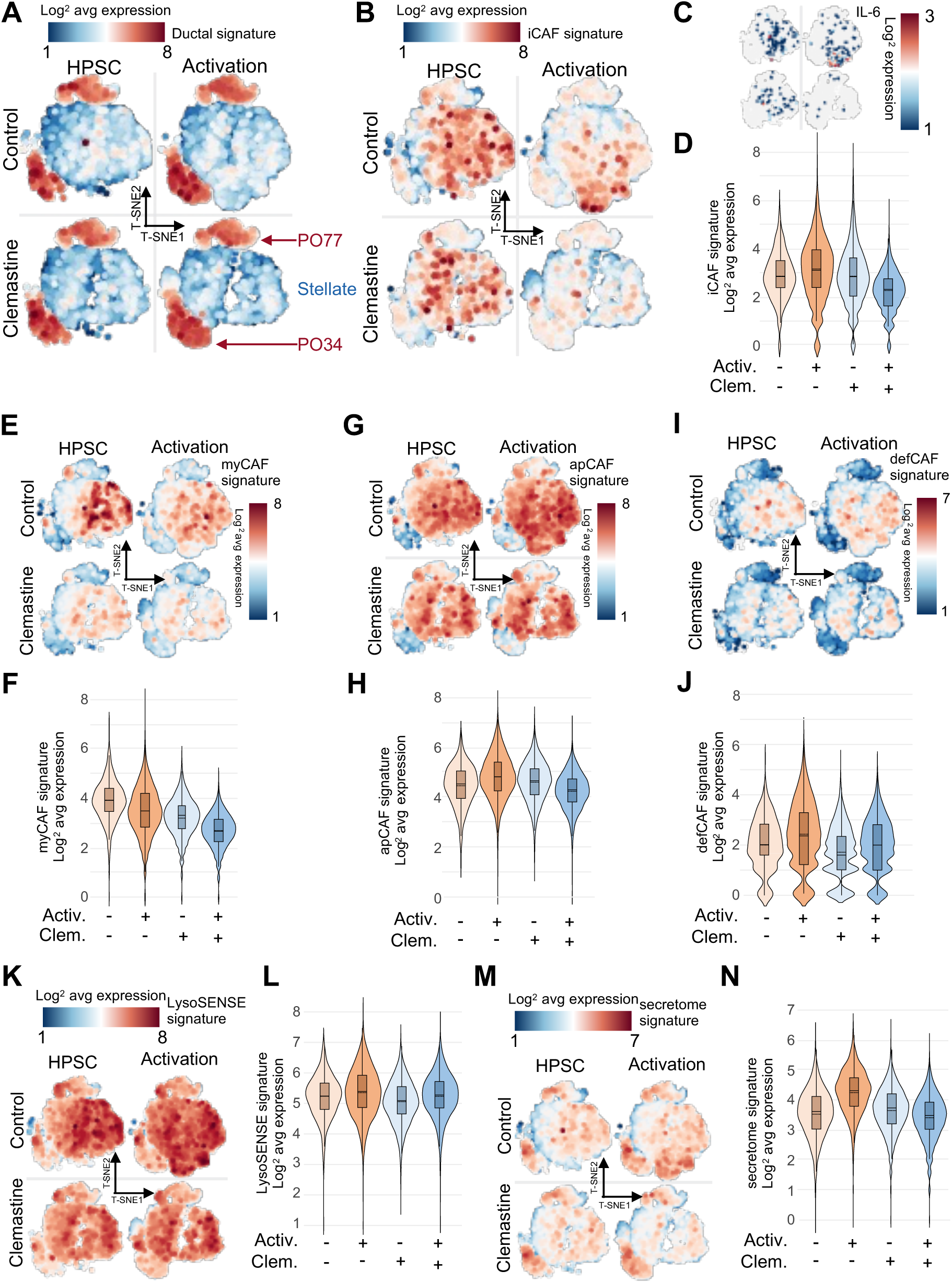
CAF identity dynamics in response to activation by serum-starvation and clemastine treatment. **A.** Integrated T-distributed Stochastic Neighbor Embedding (T-SNE) plot of 49210 barcoded cells including parental and activated HPSCs alone or in Matrigel co-cultures with PO34 and PO77, in control and clemastine-treated conditions (4 µg/mL, 24h). Gene expression signature for neoplastic ductal cells (as per Fig. 7C-E) highlights PO34 and PO77 cells (red) from stellate (blue). **B**. T-SNE plot of parental and serum-starved activated HPSCs alone or in Matrigel co-cultures with PO34 and PO77, in control and clemastine-treated conditions (4 µg/mL, 24h). Gene expression signature for putative iCAFs (expressed as log2 average gene expression from blue to red). **C**. T-SNE plot of parental and activated HPSCs alone or in Matrigel co-cultures with PO34 and PO77, in control and clemastine-treated conditions (4 µg/mL, 24h). Gene expression signature for IL-6 (expressed as log2 gene expression from blue to red). **D.** Violin plot for the gene expression distribution for putative iCAFs (expressed as log2 average gene expression) in parental or serum-starved activated HPSCs, in control and clemastine-treated conditions (4 µg/mL, 24h). **E**. T-SNE plot of parental and serum-starved activated HPSCs alone or in Matrigel co-cultures with PO34 and PO77, in control and clemastine-treated conditions (4 µg/mL, 24h). Gene expression signature for myelofibroblastic myCAFs (expressed as log2 average gene expression from blue to red). **F.** Violin plot for the gene expression distribution for myCAFs (expressed as log2 average gene expression) in parental or activated HPSCs, in control and clemastine-treated conditions (4 µg/mL, 24h). **G**. T-SNE plot of parental and serum-starved activated HPSCs alone or in Matrigel co-cultures with PO34 and PO77, in control and clemastine-treated conditions (4 µg/mL, 24h). Gene expression signature for antigen presenting apCAFs (expressed as log2 average gene expression from blue to red). **H.** Violin plot for the gene expression distribution for apCAFs (expressed as log2 average gene expression) in parental or serum-starved activated HPSCs, in control and clemastine-treated conditions (4 µg/mL, 24h). **I**. T-SNE plot of parental and serum-starved activated HPSCs alone or in Matrigel co-cultures with PO34 and PO77, in control and clemastine-treated conditions (4 µg/mL, 24h). Gene expression signature for defender/tumor suppressive defCAFs (expressed as log2 average gene expression from blue to red). **J.** Violin plot for the gene expression distribution for defCAFs (expressed as log2 average gene expression) in parental or serum-starved activated HPSCs, in control and clemastine-treated conditions (4 µg/mL, 24h). **K**. T-SNE plot of parental and serum-starved activated HPSCs alone or in Matrigel co-cultures with PO34 and PO77, in control and clemastine-treated conditions (4 µg/mL, 24h). Gene expression signature for LysoSENSE (expressed as log2 average gene expression from blue to red). **L.** Violin plot for the gene expression distribution for LysoSENSE (expressed as log2 average gene expression) in parental or serum-starved activated HPSCs, in control and clemastine-treated conditions (4 µg/mL, 24h). **M**. T-SNE plot of parental and serum-starved activated HPSCs alone or in Matrigel co-cultures with PO34 and PO77, in control and clemastine-treated conditions (4 µg/mL, 24h). Gene expression signature for secretome markers (as per Fig. 6) (expressed as log2 average gene expression from blue to red). **N.** Violin plot for the gene expression distribution for secretome markers (expressed as log2 average gene expression) in parental or activated HPSCs, in control and clemastine-treated conditions (4 µg/mL, 24h).

Lysosomal damage sensing can be estimated by the expression of lysosome-specific genes, including *LGALSs*, *LAMPs*, and other regulators of biological membrane homeostasis (**Supplementary Fig. 7E**). We defined this gene signature as LysoSENSE and investigated its enrichment in HPSCs and PDAC cells following HPSC activation and/or clemastine exposure (**Fig. 8K-L**). While LysoSENSE signature score remained stable in both PO34 and PO77 PDAC cells, it was specifically upregulated in activated HPSCs (**Fig. 8K-L**) indicating a CAF-specific lysosomal stress sensing. Clemastine treatment led to a slight reduction in both LysoSENSE signature score and cell abundance, likely reflecting its cytotoxicity on iCAF- and myCAF-like HPSCs. Low LysoSENSE score and absence of change in response to clemastine treatment in PO34 and PO77 further support the *in vitro* and *in vivo* data across the entire study, indicating that clemastine’s cytotoxic effects are preferentially directed towards CAFs rather than PDAC cells.

To complete our analysis of the PDAC microenvironmental features, we studied gene expression enrichment for cytokines that showed significantly increased abundance in patient avatar samples treated with clemastine (**Fig. 6**). We confirmed that the expression levels of these cytokines increased in activated iCAF-like HPSCs but globally decreased following clemastine exposure (**Fig. 8M-N**).

### Clemastine decreases disease burden and halts metastatic progression

As clemastine reduced the tumor burden in the PDAC patient avatars by primarily attacking the CAF populations, we next assessed whether clemastine treatment affected the metastatic dissemination of the tumors. At 6-8 weeks post-implantation, vehicle-treated patient avatars from both PO34 and PO77 exhibited macroscopic signs of pancreatitis and edema, the latter suggestive of liver metastases and/or lymph node dissemination (**Fig. 9A**). To analyze possible metastatic spread, were performed histopathological quantification of metastases in the spleen, liver, and lymph nodes using antibodies specific to the human nuclear antigen (NUMA) to visualize tumor cells. In the vehicle groups, metastatic dissemination was observed in 100% of PO34 and 83% of PO77 patient avatars in at least one of the organs (**Fig. 9B-D**). In most of the PO34 and all PO77 avatars clemastine monotherapy effectively limited metastatic spread as only 29% and 0% of avatars showed metastases, respectively. This was comparable to the gemcitabine monotherapy (17% vs 0%, respectively). Strikingly, no metastases were found in the combination avatars of the PO34 model (**Fig. 9B**), suggesting synergistic effects of clemastine and gemcitabine in inhibition the metastatic dissemination.

**Figure 9.**
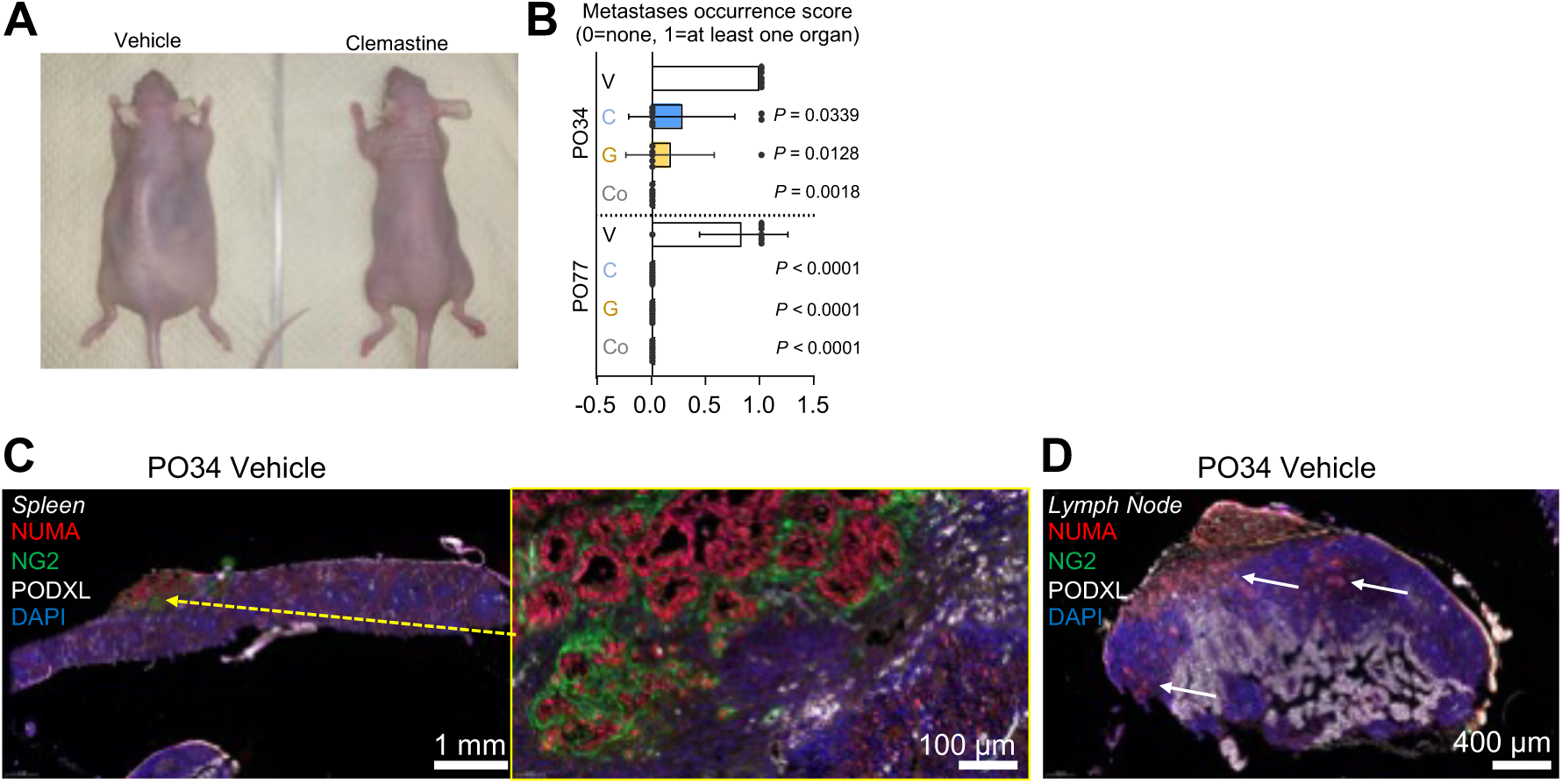
Clemastine improves patient avatar prognosis by reducing metastatic progression in vivo. **A**. Typical pancreatitis and metastasis-associated edema in the vehicle-treated cohort (left) occurring between 6-8 weeks that was not observed in the clemastine-treated group (right). **B.** Metastasis occurrence scoring (0=none detected, 1=at least one organ) in the spleen, liver and lymph nodes. Pathology performed in PO34 PDAC patient avatars from the vehicle (V, white, N=5), clemastine (C, blue, 50 mg/kg, N=7), gemcitabine (G, yellow, 100 mg/kg, N=7), and combination (Co, grey, N=8) as well as from PO77 PDAC patient avatars from the vehicle (V, white, N=6), clemastine (C, blue, 50 mg/kg, N=9), gemcitabine (G, yellow, 100 mg/kg, N=9), and combination (Co, grey, N=10) cohorts. *P*-values calculated with Kruskal-Wallis multiple comparison tests to the Vehicle control. **C.** Representative micrograph of the metastatic dissemination of PO34 PDAC cells into the spleen. Human tumor cells (NUMA, red), cancer-associated fibroblasts (NG2, green), blood vessels (podocalyxin, PODXL, white), and cell nuclei (DAPI, blue). Middle panel shows higher magnification of the metastatic lesion from the right panel marked with yellow arrow. **D.** Representative micrograph of the metastatic dissemination of PO34 PDAC cells into the neck lymph node. Human tumor cells (NUMA, red), cancer-associated fibroblasts (NG2, green), blood vessels (PODXL, white), and DAPI (blue). NG2=chondroitin sulphate proteoglycan 4, PODXL=podocalyxin.

## Discussion

PDAC stroma and microenvironment composition are essential to support the neoplastic cells and fuel cancer progression and metastasis. In this study, we found that half of the tested PDAC POs showed *in vitro* sensitivity (i.e decreased cancer cell viability) to clemastine, a cationic amphiphilic drug that induces lysosomal membrane permeabilization and cell death. Interestingly, the *in vitro* sensitivity profiles did not predict *in vivo* response, illustrating a limitation of cellular models for preclinical research and precision medicine applications. Instead, clemastine unexpectedly led to a dramatic depletion of CAFs, halted tumor progression, and inhibited metastasis *in vivo* in patient avatar models, regardless of their original *in vitro* sensitivity to clemastine.

Therapeutic targeting of CAFs in PDAC has been the focus of many recent studies. However, due to the diversity of CAFs, development of treatments targeting CAFs has been challenging ^7,8^. It has also become clear that the tumor-TME interactions are not uniformly tumor promoting or suppressive, and the underlying mechanisms remain incompletely understood.

iCAF and myCAF subtypes are dynamically interchangeable and attempts to exclusively deplete αSMA^+^ fibroblasts (myCAF) led to shortened survival in preclinical studie.s^37^ A cellular shift between myCAF and iCAF subtypes enabled PDAC cells to migrate away from their protective periglandular myCAF niche, while the expansion of the iCAF population further promoted tumor immunosuppression.^8^ Ongoing trials should therefore aim at blocking CAF plasticity and/or eradicate all CAF subtypes. Our data show that clemastine acted as an all-in-one CAF destructor, selectively targeting tumor supporting iCAFs and myCAFs. Since clemastine showed limited efficacy on PDAC cells in vivo, it holds great repurposing potential as a companion therapy in precision medicine, as we demonstrate in combination with gemcitabine.

Lysosomes are highly dynamic organelles central to degradation and recycling of molecules, and their pivotal role in nutrient sensing has recently gained a lot of interest.^38,39^ In addition to their cancer promoting functions^14^, lysosomes in cancer cells are particularly sensitive to _LMP.40-42_

Our results show that CAFs are particularly susceptible to CAD-induced LMP. PDAC cells have been shown to switch energy source from glucose and serum-derived nutrients to pancreatic stellate cell (PSC)-derived alanine to drive the tricarboxylic acid cycle. This alanine secretion is dependent on the PSC lysosomal activity and stimulated by cancer cells.^43^ This supports the idea of increased lysosomal dependence during malignant transformation and subsequently increased sensitivity to lysosomal damage of CAFs. In cancer and other fibrotic diseases, normal fibroblasts exposed to exosomes derived from activated myofibroblast convert into myofibroblasts, acquiring enhanced proliferative and migratory properties.^44^

Here, we derived iCAF-like cells from fibroblast precursors, HPSCs, by serum-starvation. This mimics the nutrient-poor milieu of tumor where the (i)CAFs must survive. The starvation caused lysosomal stress (LGALS1 puncta), upregulated the lysosomal gene signature, LysoSENSE, and increased clemastine sensitivity. When exposed to the lysosomal-targeting effects of clemastine, starved HPSCs (iCAF-like cells) – as well as the CAFs in the patient avatars – were more sensitive to cell death than their healthy counterparts.

It has been shown that PDAC CAFs are intrinsically resistant to gemcitabine and gemcitabine-exposed CAFs promote PDAC cell proliferation and drug resistance.^45^ Thus, CAF-provided support to cancer cells not killed by gemcitabine would continue even after the treatment. In addition, off-targeting has been a fundamental issue in trials focusing on depleting CAFs.^46^ Here we demonstrate a clear advantage of clemastine over current CAF-targeting therapies by targeting the CAF vulnerability to LMP not shared by the healthy pancreatic fibroblasts. Clemastine also potently reshaped the tumor-supporting stroma into a microenvironment that further increased efficacy of conventional therapies with on cancer cells, as we report for gemcitabine.

We detected no *in vitro* synergy between clemastine and other tested chemotherapy regimens in vitro. However, gemcitabine, along with other classical chemotherapy drugs, is known to cause a chemo-induced fibrotic reaction^47^. This reaction hampers the influx of further cytostatic compounds and leads to an acquired physical chemoresistance. A fibrotic reaction was, indeed, detected in our patient avatars upon gemcitabine treatment, and reversed by the addition of clemastine. This proposes that while no direct synergy is seen between these compounds, clemastine could inhibit or postpone the development of the fibrotic reaction and thus prolong the effectiveness of gemcitabine (and perhaps other regimens). Further studies are however needed to test this hypothesis. Another evident synergistic advantage of the combination treatment with clemastine and gemcitabine was the complete inhibition of metastasis.

The main strength of our study is the layering of in vitro and in vivo findings as well as integration of functional drug testing with both proteomic and transcriptomic data. There are, however, also weaknesses in the study. The starvation of HPSCs was performed to induce cancer cell activating, iCAF-like phenotype. The deprivation from nutrients does mimic the nutrient-poor tumor environment of the PDAC CAFs. Serum does, however, contain more than just nutrients, and the changes in transcriptomics occurring in the serum-starved HPSCs may be affected by lack of the many cytokines that are present in sera. Our results do, however, demonstrate that starvation induced the iCAF gene signature, which was the principal aim of the starvation, and promoted sensitivity to clemastine.

Clemastine has potential to be used in combination with gemcitabine and other therapeutic regimens in PDAC. As clemastine is well-tolerated and does not hamper wound healing, it can further be administered as a single agent at times when conventional regimens cannot be given – such as pre- and post-surgery. Being well-tolerated, clemastine could also be used as maintenance treatment during the follow-up after the final cycles of adjuvant. Thus, we propose clemastine as a potential therapeutic in PDAC.

## Materials and Methods

### Cell cultures

We used three commercially available PDAC 2D cell lines for our experiments: MiaPaCa-2, a primary tumour cell line, and two metastatic tumour cell lines: HPAF-II and AsPC-1. The cell lines were purchased from the American Type Culture Collection (ATCC) and cultured according to the provider’s instructions.

The human skin fibroblast cell line WS1 (ATCC) was cultured according to the provider’s instructions. The immortalised human pancreatic stellate cell line, HPSC, kindly provided by Dr. Hwang, MD Anderson Cancer Center, Houston, TX, USA^33^ was cultured in DMEM supplemented with 10% FBS. To activate the HPSC cells prior treatment, they were serum-starved for 3 days.

### Patients and sample processing

The study setting constituted of four patients operated at the Helsinki University Hospital between 2018 and 2019. Only patients with confirmed PDAC were included in the study. Median age at operation was 68 (63-73) years (Fig. 2A). Patient records and the Finnish Population registry were used to obtain the clinical information. Patients were followed up for a median 2.8 years (range 1.6 to 4.6). Follow-up ended at death or in April 2024. Three out of four patients had succumbed to the disease at the end of the follow-up period, while one patient (PO80) was still alive. The study was approved by the University of Helsinki ethical committee, and the pancreatic cancer research protocol (HUS 2075/2016) was applied. Written informed consent was received from all patients.

The freshly resected surgical pancreatic specimen was transferred as such to the department of pathology, where it was examined and opened by an experienced gastrointestinal pathologist. The tumour was macroscopically located and an approximately 15 mm x 5 mm x 2 mm tumour sample was transferred to the laboratory for organoid culture (transported on ice in human wash medium. ^42^ A corresponding piece of surrounding tissue was cut for histology in a separate cassette to ascertain the malignancy of the specimen.

The resected tumour tissues were processed according to the Tuveson laboratory protocol.^42^ In short, the sample was digested first mechanically and then chemically in human digestion medium (see below) at 37°C for 45 min. After this, the established single cell suspension was washed twice in human wash medium before plating the cells in Corning® Matrigel® Matrix (Fisher Scientific) domes and cultured in human complete feeding medium. The human complete feeding medium additionally supplemented with 1mM Prostaglandin E2 (PGE2) was used for growing organoids from the tumour adjacent healthy pancreatic tissue.

- Human Wash Media: Advanced DMEM/F-12 supplemented with 1% GlutaMAX, 10 mM HEPES, 100 mg/ml Primocin, and 0.1% BSA
- Human Complete Feeding media: [Human Wash Media] supplemented with 50% Wnt3a-Conditioned Medium, 10% R-spondin1-Conditioned medium, 1% B27 Supplement, 10 mM Nicotinamide, 1.25 mM N-acetylcysteine, 100 ng/mL mNoggin, 100 ng/ml hFGF, 50 ng/mL hEGF, 10 nM hGastrin-I, 500 nM ALK4, ALK5 and ALK7 inhibitorA83-01
- Human Digestion Media: [Human Complete Feeding media] supplemented with 10.5 mM ROCK inhibitor Y-27632, 5mg/ml Collagenase XI, and 10mg/ml DNAse I

### DNA sequencing

Next-generation sequencing of DNA extracted from the primary surgical tissue as well as from the established patient derived organoids (POs) and 2D patient derived cells was performed at FIMM Technology Center (using Comprehensive Cancer Panel by NimbleGen) for PO34, PO34A, and PO37. EDTA-blood samples were also collected preoperatively from the patients. DNA was isolated from the blood and was used as germline control for sequencing at FIMM. As the use of NimbleGen Comprehensive Cancer Panel at FIMM stopped, the following samples (PO77, PO80) were sequenced at HUSlab (using Ion AmpliSeq Cancer Hotspot Panel v2 by ThermoFisher).

### Patient derived cell cultures

Organoids were passaged by suspending the Matrigel domes containing the cells in Corning® Cell Recovery Solution (Fisher Scientific). After a 30-minute incubation on ice, the organoids were treated with TrypLe™ (Gibco™) for 5 min in 37°C, washed twice with human wash medium and plated again in Matrigel® domes. Following each passage of organoids, the human complete feeding medium was freshly supplemented with 10.5 mM ROCK inhibitor Y-27632.

The fibroblast cell line CAF82 was established in our laboratory. The fibroblasts started growing out from the PDAC organoid cultures and were left attached while organoids were detached upon passaging. Dulbecco’s Modified Eagle’s Medium (DMEM) supplemented with 10% FBS and 100 µg/ml Primocin was used for the culture of the fibroblasts. Once the fibroblasts grew to 70% confluency, they were passaged conventionally by using trypsin to detach the cells.

### In vitro drug sensitivity testing

We used a 96-well plate to study the sensitivity of the commercial PDAC cell lines, fibroblasts, and POs to clemastine. For this, 3000-5000 cells/well were plated (according to their growth rate) either directly on plastic (the 2D cell lines) or suspended in 5 µL Matrigel (organoids), and 100 µL medium was added to the culture. Before plating the organoids, they were dissociated into single cells using the Tuveson protocol ^42^. In short, the organoid-containing Matrigel domes were suspended in human wash medium supplemented with 2 mg/mL Dispase II and 10 ug/mL DNAse-I and incubated in 37°C for 20 min. After this, the domes were collected, and the suspension was examined under a light microscope. If the organoids had not been dissolved into a single-cell solution, an additional 5-minute trypsinization at 37°C was performed.

The day after seeding, clemastine (in increasing concentrations, Supplementary Table S1). DMSO served as a negative and Benzethonium chloride (BzCl), a toxic compound, ^48^ was used as a positive control. After a 3–6-day incubation, the cell viability was assessed either with CellTiter-Glo reagent (CTG, Promega, Madison, WI), a luminescence -based assay, or using 10 μL of 3-(4,5-dimethylthiazol-2-yl)-2,5-diphenyltetrazolium bromide (MTT; 5 mg/ml in PBS) incubated for 2h at 37°C. After MTT treatment, cells were lysed in 10% SDS-10 mM HCl overnight and absorbance was measured at 540 nm using Multiskan Ascent software version 2.6 (Thermo Labsystems). The experiments were repeated three times.

The IC50, EC50, and DSS as well as quality control for the screens were found for the Individual drugs from the raw data using a FIMM in-house analytics tool Breeze (https://breeze.fimm.fi/).^43^ A DSS >10 was considered sensitive.

To study whether clemastine synergized with other drugs commonly used in treatment of PDAC, we seeded the organoid-derived single-cell suspension in a 50% Matrigel (stock 8–10 mg/mL) and 50 % human complete feeding medium mixture in 384-well plates (Corning). The final concentration of cells was 5000–10 000 per 10 μL dome. Human complete feeding medium was used as growth medium. Once organoids started to form – usually 2-3 days after seeding – three approved regimens (Gemcitabine, Paclitaxel and FOLFIRINOX) and one investigational oncology drug (Trametinib) were added to the culture at increasing concentrations (Supplementary Table S1). The drugs were used both alone and in combination with a fixed concentration (14 nM) of clemastine to study if any synergies could be found. FOLFIRINOX is a combination of four drugs - Folinic acid (also called leucovorin), fluorouracil (5-FU), irinotecan (or its active metabolite SN-38), and Oxaliplatin. The ratio between the drug concentrations in FOLFIRINOX can be seen in Supplementary Table S1. After a 6-day incubation, cell viability was assessed by using CTG, and results were analysed with the Breeze analytics tool.

The occurrence of lysosomal membrane permeabilization (LMP) in clemastine-treated and control cells was measured using an overnight 2 µg/ml clemastine concentration followed by visualization of LAMP2 and galectin-1 (LGALS1). Cells were fixed with 4% PFA for 10 min at RT, followed by washes with PBS and permeabilization with 0,3% Triton X-100 (Sigma) for 5 min at RT. Primary antibodies against LAMP2 (mouse anti-LAMP2, 1:200) and LGALS1 (rabbit anti-LGALS1, 1:100) were incubated with fixed cells for 60 min RT followed by 3x5 min washes with PBS. Secondary antibodies donkey anti-mouse Alexa-488 (1:1000, Invitrogen) and donkey anti-rabbit Alexa-568 (1:1000, Invitrogen), respectively were incubated for 30 min at RT followed by 1xwash with PBS. Cell nuclei were visualized by using Hoechst (Invitrogen) staining.

### Cytokine array

Secretion of selected human cytokines and chemokines to the cell culture medium was determined using the proteome profiler human or mouse XL cytokine array kits (R&D Systems). Briefly, 500 μL of conditioned media from control and clemastine-treated CAF82 and PO34 cell cultures or whole tissue extracts pooled from 3 independent PO77 PDAC patient avatars from the vehicle, clemastine, gemcitabine or combination treatment groups were run on the array according to the manufacturer’s instructions. Signal was visualized using chemiluminescent detection and developed on X-ray film. The average pixel density of each analyte from duplicate spots was determined and corrected according to reference dots. To determine the influence of chemotherapies, dot intensity ratio was calculated as ratio to the vehicle control.

### Patient avatars

Animal experiments were approved by the Committee for Animal Experiments of the District of Southern Finland (ESAVI/6285/04.10.07/2014 and ESAVI/10262/2022). Six-weeks-old immunocompromised female NMRI-nu (Rj:NMRI-Foxn1^nu^/Foxn1^nu^, Janvier Labs) mice were anesthetized using a Ketamine/Xylazine cocktail and placed on a heating pad. The left side of the animal’s flank, below the thorax, was disinfected using a Biseptine solution. Temgesic (0,12 mg/kg in 100 µL saline) was injected subcutaneously next to the spleen. A six mm incision of the skin was performed with small, curved scissors to access the muscle layer. A four mm incision was performed with sharp surgery scissors to access the spleen. The spleen was then carefully pulled out of the peritoneum with forceps and 5 mL of cell suspension (50000 cells) injected in the splenic vein upstream from the mouse pancreas using a 30G Myjector single use insulin syringe. The needle was carefully removed, and a piece of sterile absorption triangle (Fine Science Tools, Germany) was applied to stop the bleeding when occurring. Spleen was then gently placed back into the peritoneal cavity. Muscle layer was sutured first, separately followed by the skin. Biseptine was sprayed on the wound and the animal placed back into the cage and monitored until full recovery. Temgesic (0,12 mg/kg in 100 µL saline) was injected subcutaneously next to the wound daily for the three following days. Twenty days post-surgery, chemotherapeutic regimen was initiated according to the following section’s protocol.

### Chemotherapy of the patient avatars

#### Gemcitabine

PO34 patient avatars were carefully maintained on the back, with the index finger behind the head to ensure that the head was not moving sideways when inserting the needle. A flexible single use polypropylene 20G feeding needle was gently inserted at the left or right edge of the mouse’s mouth and pushed vertically in the trachea, following the curvature of the neck. When 30 mm was inserted, 100 µL of gemcitabine (100 mg/kg, in saline) was delivered and the tube carefully retracted. The mouse was then placed back in the cage and its behaviour (locomotion, breathing) monitored during the following ten minutes. Gemcitabine was administered every three days for up to fifty days.

PO77 patient avatars received gemcitabine (100 mg/kg, in saline, 3 µL/s) as an intraperitoneal injection, using a 30G Myjector single use insulin syringe, every three days for up to forty-five days.

#### Clemastine

as previously described^16^, both PO34 and PO77 patient avatars received clemastine dissolved in saline as an intraperitoneal injection, using a 30G Myjector single use 300 µL insulin syringe, at an initial 100 mg/kg (in 200 µL saline, 3 µL/s) on day one, followed by daily intraperitoneal injections at 50 mg/kg (100 µL, in saline, 3µL/s).

#### Combination

PO34 and PO77 received oral dosage of gemcitabine and intraperitoneal injection of clemastine as described above. On combination treatment days, two hours between the gemcitabine oral/peritoneal dosage and the clemastine injection were given to prevent potential undesired drug interactions.

### Antibodies and reagents

For the immunofluorescence detection patient avatar tissue sections were stained by using rat monoclonal anti-mouse podocalyxin (PODXL,MAB1556, Cell Signalling, dilution 1:800), rabbit polyclonal anti-NG2 (AB5320, R&D Systems, dilution 1:200), mouse monoclonal Cy3-conjugated anti-human nuclear mitotic apparatus protein (NUMA, MAB1281C3, Cell Signalling, dilution 1:400), mouse monoclonal Cy3-conjugated anti-α-SMA (C6198, Sigma-Aldrich, dilution 1:200), rabbit polyclonal anti-mouse PDGFR-β (sc-436, Santa Cruz Biotechnology, dilution 1:50), mouse monoclonal anti-HCAM (sc-7297, Santa Cruz Biotechnology, dilution 1:200), rabbit polyclonal antibodies against LGALS1 (ab25138, Abcam Biotechnologies, dilution 1:600) and LAMP2 (ab25631, Abcam Biotechnologies, dilution 1:200). DAPI (cat. 5748, Tocris, 1µg/mL) was used to visualize the cell nuclei.

The in vitro HPSC cultures were stained by using monoclonal mouse anti-FAP (66562, Cell Signalling, dilution 1:100), Vimentin (sc-6260, Santa Cruz Biotechnology, dilution 1:500), or aSMA (MAB1420, R&D Systems, dilution 1:2500). Donkey anti-mouse (AlexaFluor488, A21202, Invitrogen, dilution 1:1000) and donkey anti-rabbit (AlexaFluor488, A21206, Invitrogen, dilution 1:1000) were used as secondary antibodies. Hoechst 33342 10mg/ml (H3570, Invitrogen, dilution 1:5000) was used as a nuclear counterstain.

### Immunofluorescence staining of xenograft tumour tissue

Snap-frozen tissues (pancreas, lymph nodes, liver, spleen) were cut using a cryotome (Leica CM3050) into 9-μm-thick coronal sections serried on microscope slide (5-10 sections per slide, 15 slides total). Prior to immunofluorescence/histochemistry, tissue sections were fixed in 4% PFA, blocked with 5% FBS (Lonza) and 0.03% Triton X-100 (Sigma) in PBS. Whole slides were scanned using a slide scanner (3DHISTECH Panoramic P250 FLASH II digital slide scanner). Tumour volumes were calculated using the histological coordinates defined by the tissue section series. If antibody species were incompatible for simultaneous co-staining, verification was performed using consecutive microscope slides, on 9-μm distant consecutive tissue sections. Tissue structures (e.g. blood vessels, CAFs, etc.) were used as spatial references. Histology quantification (e.g., tumour cell invasion, blood vessel density, pericytes coverage, etc.) was performed on 10 sections equally distributed along the entire tissue, for each patient avatar. For statistical analyses, individual data points are averaged values for all counted sections.

### Microscopic imaging

For the transmitted light microscopy, the cells were visualized and imaged using an inverted epifluorescence microscope (Zeiss Axiovert 200) equipped with AxioVision software. Confocal images were taken with the Zeiss LSM 780 and 880 equipped with appropriate lasers. HPSC cultures were visualised using the 40x Olympus PlanSApo objective. MicroManager 2.0 software was used to take the images, which were then processed with ImageJ.

For the patient avatar histological analyses, Pannoramic viewer (3DHISTECH) annotation and quantification tools were used to perform total tumour and necrotic area quantification. Metastatic lesions were counted manually in selected organs. Automated cell counting, co-staining and stained surface calculation were performed in Fiji (https://doi.org/10.1038/nmeth.2019) using the *Cell Counter*, *Analyse Particles*, *2D Histogram* and *Volume Calculator* embedded plugins, after transformation of individual composite pictures into binary images using the *make binary* function.

### Single cell sequencing (scRNAseq)

Clemastine treatment induced ranscriptomic changes in POs and HPSCs were assessed using scRNAseq. First, monocultures of HPSCs and co-cultures of POs and HPSCs (ratio 1:2) were seeded within Matrigel/Collagel (4.35 mg/mL:0.5 mg/mL) mixture domes on 24-well corning plates. One half of the cultures were grown in Advanced DMEM: f12+Rspo 10% + WNT 50% media containing 10% FBS. The other half of the cultures were grown in Advanced DMEM: f12+Rspo 10% + WNT 50% serum-free media for HSPC activation. After a 24h establishment of the mono-/co-cultures, clemastine (4 µg/mL)/DMSO control was added.

To analyse the transcriptomic alterations occurring before the eventual cell death, clemastine-treatment was stopped after 24 hours. The domes were collected, and organoids dissolved into a single-cell solution according to above-described method.

After gaining the single-cell solution, we performed cell hashing ^49^ using four different hashtags, enabling us to pool cell cultures that had been treated according to the same schemes; starved/non-starved hPSC samples were pooled together as were the clemastine-treated/untreated samples. Cell hashing analysis was performed according to the TotalSeq™-A Antibodies and Cell Hashing with 10x Single Cell 3’ Reagent Kit v3.1 (Dual Index) Protocol (https://www.biolegend.com/en-us/protocols/totalseq-a-dual-index-protocol). Count matrix for Hashtag oligonucleotides (HTO) were further generated using the CITE-seq-Count-tool (https://zenodo.org/records/2590196).

Finally, scRNA expression profiles were analysed by using the 10x Genomics Chromium Single Cell 3’RNAseq platform. Chromium Next GEM Single Cell 3’ Gene Expression version 3.1 Dual Index chemistry was used for the Chromium Single Cell 3’RNAseq run and library preparation. Sample libraries were sequenced on Illumina NovaSeq 6000 system using read lengths: 28bp (Read 1), 10bp (i7 Index), 10bp (i5 Index) and 90bp (Read 2). Data processing and analysis were performed using the 10x Genomics Cell Ranger v7.1.0 pipelines (“cellranger mkfastq” to produce FASTQ files and “cellranger count” for alignment, filtering and UMI counting). GRCh38 was used as reference genome for alignment and mkfastq was run using the Illumina bcl2fastq v2.2.0. Further, the cellranger aggr pipeline was used to aggregate individual samples into a single feature-barcode matrix.

R (version 4.2.3, Foundation for Statistical Computing, Vienna, Austria) with packages Seurat (v5.01), Celldex (v.1.8.0), SingleR (v2.0.0), EnhancedVolcano (v. 1.16.0), DoubletFinder (c2.04), scRNAseq (v.2.12.0), Scuttle (v. 1.8.4) were used for generating UMAPs and analysis of gene deregulation. R-studio (v. 2023.06.1) Posit software, Boston, MA) was the integrated development environment (IDE) used. Raw data was filtered to include only high-quality cells defined by detectable genes between 250-7000, log10(genes/unique molecular identifier (UMI)) >0.8 and mitochondrial genes <15%. Data was demultiplexed with HTODemux from the Seurat package, removing positive cells for more than one hashtag oligo (HTO) or negative to all HTOs. Doublet finder was applied for additional cell filtering, and filtered data were log normalized, scaled with regressing out cell-cell variation due to UMI counts, percent mitochondrial reads and cell cycle genes. We annotated the cells with the SingleR package using celldex human pancreas scRNA experiment^44^ as a reference database. Integration of data was performed with Seurat’s reciprocal principal component analysis (RPCA) with 2000 variable features and 15 dimensions. Further dimensional reduction was obtained by using Uniform Manifold Approximation and Projection (UMAP).

### Gene set enrichment analysis

Gene set enrichment analysis was performed using gseGO function in the clusterProfiler (v. 4.6.2) R package with org.Hs.eg.db (v. 3.16.0) database for genome wide annotation for human genes and biological processes as Go aspect. Genes were ranked with average log2(fold control) values obtained from differential gene expression results.

### Statistical analysis

All statistics were computed using the GraphPad Prism 9 (GraphPad, La Jolla, CA, USA) and IBM software SPSS Statistics for Windows, version 22 (IBM Corp., Armonk, NY, USA). Statistical significance between sample groups was determined by *P*-values calculated by unpaired two-tailed t-test or the non-parametric Mann-Whitney *U*-test. For multiple comparisons, one-way ANOVA with the Kruskal-Wallis or Holm-Sidak’s multiple comparisons tests were used. Data are expressed as mean ± SD or ± SEM and considered as statistically significant when P < 0.05. Statistical tests are specified in the figure legends. All images and graphs shown are representative of several experiments as indicated in the figure legends.

## Acknowledgements

We acknowledge Henrikki Almusa (FIMM, UH) for exome sequencing analysis, FIMM Single-Cell Analytics and FIMM Genomics Sequencing units supported by HiLIFE and Biocenter Finland for single cell sequencing services, Biomedicum Imaging Unit for their assistance in immunofluorescence microscopy imaging, Genome Biology Unit (Faculty of Medicine, University of Helsinki, Biocenter Finland) for the 3DHISTECH Pannoramic P250 FLASH II digital slide scanner service. We also acknowledge the HiLIFE Laboratory Animal Centre Core Facility, University of Helsinki, Finland for the support with animal experimentation. Further, we express our gratitude to the Juselius Foundation, the South-Carelian Society of Doctors and Duodecim of Viborg Lappeenranta, Finska Läkare Sällskapet, Mary and Georh Ehrnrooths foundation for supporting this work. VLJ acknowledges the support from the Magnus Ehrnrooth foundation, Academy of Finland (grant #321867) and Finnish Cancer Societies. Lastly, we want to express our gratitude to the patients who participated in the study.

## REFERENCES

1. Aaltonen, P., Carpen, O., Mustonen, H., Puolakkainen, P., Haglund, C., Peltola, K., and Seppanen, H. (2022). Long-term nationwide trends in the treatment of and outcomes among pancreatic cancer patients. Eur J Surg Oncol 48, 1087–1092. 10.1016/j.ejso.2021.11.116.

2. Allemani, C., Matsuda, T., Di Carlo, V., Harewood, R., Matz, M., Niksic, M., Bonaventure, A., Valkov, M., Johnson, C.J., Esteve, J., et al. (2018). Global surveillance of trends in cancer survival 2000-14 (CONCORD-3): analysis of individual records for 37 513 025 patients diagnosed with one of 18 cancers from 322 population-based registries in 71 countries. Lancet 391, 1023–1075. 10.1016/S0140-6736(17)33326-3.

3. Itoh, Y., Takehara, Y., Kawase, T., Terashima, K., Ohkawa, Y., Hirose, Y., Koda, A., Hyodo, N., Ushio, T., Hirai, Y., et al. (2016). Feasibility of magnetic resonance elastography for the pancreas at 3T. J Magn Reson Imaging 43, 384–390. 10.1002/jmri.24995.

4. LeSavage, B.L., Zhang, D., Huerta-Lopez, C., Gilchrist, A.E., Krajina, B.A., Karlsson, K., Smith, A.R., Karagyozova, K., Klett, K.C., Huang, M.S., et al. (2024). Engineered matrices reveal stiffness-mediated chemoresistance in patient-derived pancreatic cancer organoids. Nat Mater 23, 1138–1149. 10.1038/s41563-024-01908-x.

5. Wirth, M., and Schneider, G. (2024). A Hypoxia-Epigenetics Axis Drives EMT in Pancreatic Cancer. Cancer Res 84, 1739–1741. 10.1158/0008-5472.CAN-23-3578.

6. Grunwald, B.T., Devisme, A., Andrieux, G., Vyas, F., Aliar, K., McCloskey, C.W., Macklin, A., Jang, G.H., Denroche, R., Romero, J.M., et al. (2021). Spatially confined sub-tumor microenvironments in pancreatic cancer. Cell 184, 5577–5592 e5518. 10.1016/j.cell.2021.09.022.

7. Zhang, T., Ren, Y., Yang, P., Wang, J., and Zhou, H. (2022). Cancer-associated fibroblasts in pancreatic ductal adenocarcinoma. Cell Death Dis 13, 897. 10.1038/s41419-022-05351-1.

8. Ohlund, D., Handly-Santana, A., Biffi, G., Elyada, E., Almeida, A.S., Ponz-Sarvise, M., Corbo, V., Oni, T.E., Hearn, S.A., Lee, E.J., et al. (2017). Distinct populations of inflammatory fibroblasts and myofibroblasts in pancreatic cancer. J Exp Med 214, 579–596. 10.1084/jem.20162024.

9. Han, C., Liu, T., and Yin, R. (2020). Biomarkers for cancer-associated fibroblasts. Biomark Res 8, 64. 10.1186/s40364-020-00245-w.

10. Yang, S., Wang, X., Contino, G., Liesa, M., Sahin, E., Ying, H., Bause, A., Li, Y., Stommel, J.M., Dell’antonio, G., et al. (2011). Pancreatic cancers require autophagy for tumor growth. Genes Dev 25, 717–729. 10.1101/gad.2016111.

11. Fennelly, C., and Amaravadi, R.K. (2017). Lysosomal Biology in Cancer. Methods Mol Biol 1594, 293–308. 10.1007/978-1-4939-6934-0_19.

12. Groth-Pedersen, L., and Jaattela, M. (2013). Combating apoptosis and multidrug resistant cancers by targeting lysosomes. Cancer Lett 332, 265–274. 10.1016/j.canlet.2010.05.021.

13. Joyce, J.A. (2005). Therapeutic targeting of the tumor microenvironment. Cancer Cell 7, 513–520. 10.1016/j.ccr.2005.05.024.

14. Mohamed, M.M., and Sloane, B.F. (2006). Cysteine cathepsins: multifunctional enzymes in cancer. Nat Rev Cancer 6, 764–775. 10.1038/nrc1949.

15. Boone, B.A., Bahary, N., Zureikat, A.H., Moser, A.J., Normolle, D.P., Wu, W.C., Singhi, A.D., Bao, P., Bartlett, D.L., Liotta, L.A., et al. (2015). Safety and Biologic Response of Pre-operative Autophagy Inhibition in Combination with Gemcitabine in Patients with Pancreatic Adenocarcinoma. Ann Surg Oncol 22, 4402–4410. 10.1245/s10434-015-4566-4.

16. Perez-Hernandez, M., Arias, A., Martinez-Garcia, D., Perez-Tomas, R., Quesada, R., and Soto-Cerrato, V. (2019). Targeting Autophagy for Cancer Treatment and Tumor Chemosensitization. Cancers (Basel) 11. 10.3390/cancers11101599.

17. Wolpin, B.M., Rubinson, D.A., Wang, X., Chan, J.A., Cleary, J.M., Enzinger, P.C., Fuchs, C.S., McCleary, N.J., Meyerhardt, J.A., Ng, K., et al. (2014). Phase II and pharmacodynamic study of autophagy inhibition using hydroxychloroquine in patients with metastatic pancreatic adenocarcinoma. Oncologist 19, 637–638. 10.1634/theoncologist.2014-0086.

18. Zeh, H.J., Bahary, N., Boone, B.A., Singhi, A.D., Miller-Ocuin, J.L., Normolle, D.P., Zureikat, A.H., Hogg, M.E., Bartlett, D.L., Lee, K.K., et al. (2020). A Randomized Phase II Preoperative Study of Autophagy Inhibition with High-Dose Hydroxychloroquine and Gemcitabine/Nab-Paclitaxel in Pancreatic Cancer Patients. Clin Cancer Res 26, 3126–3134. 10.1158/1078-0432.CCR-19-4042.

19. Marmor, M.F., Kellner, U., Lai, T.Y., Melles, R.B., Mieler, W.F., and American Academy of, O. (2016). Recommendations on Screening for Chloroquine and Hydroxychloroquine Retinopathy (2016 Revision). Ophthalmology 123, 1386–1394. 10.1016/j.ophtha.2016.01.058.

20. Gulbins, E., and Kolesnick, R.N. (2013). It takes a CAD to kill a tumor cell with a LMP. Cancer Cell 24, 279–281. 10.1016/j.ccr.2013.08.025.

21. Halliwell, W.H. (1997). Cationic amphiphilic drug-induced phospholipidosis. Toxicol Pathol 25, 53–60. 10.1177/019262339702500111.

22. Ellegaard, A.M., Dehlendorff, C., Vind, A.C., Anand, A., Cederkvist, L., Petersen, N.H.T., Nylandsted, J., Stenvang, J., Mellemgaard, A., Osterlind, K., et al. (2016). Repurposing Cationic Amphiphilic Antihistamines for Cancer Treatment. EBioMedicine 9, 130–139. 10.1016/j.ebiom.2016.06.013.

23. Le Joncour, V., Filppu, P., Hyvonen, M., Holopainen, M., Turunen, S.P., Sihto, H., Burghardt, I., Joensuu, H., Tynninen, O., Jaaskelainen, J., et al. (2019). Vulnerability of invasive glioblastoma cells to lysosomal membrane destabilization. EMBO Mol Med 11. 10.15252/emmm.201809034.

24. Makinen, L., Vaha-Koskela, M., Juusola, M., Mustonen, H., Wennerberg, K., Hagstrom, J., Puolakkainen, P., and Seppanen, H. (2022). Pancreatic Cancer Organoids in the Field of Precision Medicine: A Review of Literature and Experience on Drug Sensitivity Testing with Multiple Readouts and Synergy Scoring. Cancers (Basel) 14. 10.3390/cancers14030525.

25. Yadav, B., Pemovska, T., Szwajda, A., Kulesskiy, E., Kontro, M., Karjalainen, R., Majumder, M.M., Malani, D., Murumagi, A., Knowles, J., et al. (2014). Quantitative scoring of differential drug sensitivity for individually optimized anticancer therapies. Sci Rep 4, 5193. 10.1038/srep05193.

26. Haeberle, L., and Esposito, I. (2019). Pathology of pancreatic cancer. Transl Gastroenterol Hepatol 4, 50. 10.21037/tgh.2019.06.02.

27. Eurola, A., Ristimaki, A., Mustonen, H., Nurmi, A.M., Hagstrom, J., Haglund, C., and Seppanen, H. (2021). Impact of histological response after neoadjuvant therapy on podocalyxin as a prognostic marker in pancreatic cancer. Sci Rep 11, 9896. 10.1038/s41598-021-89134-2.

28. Saukkonen, K., Hagstrom, J., Mustonen, H., Juuti, A., Nordling, S., Fermer, C., Nilsson, O., Seppanen, H., and Haglund, C. (2015). Podocalyxin Is a Marker of Poor Prognosis in Pancreatic Ductal Adenocarcinoma. PLoS One 10, e0129012. 10.1371/journal.pone.0129012.

29. Ohlund, D., Elyada, E., and Tuveson, D. (2014). Fibroblast heterogeneity in the cancer wound. J Exp Med 211, 1503–1523. 10.1084/jem.20140692.

30. Aits, S., Kricker, J., Liu, B., Ellegaard, A.M., Hamalisto, S., Tvingsholm, S., Corcelle-Termeau, E., Hogh, S., Farkas, T., Holm Jonassen, A., et al. (2015). Sensitive detection of lysosomal membrane permeabilization by lysosomal galectin puncta assay. Autophagy 11, 1408–1424. 10.1080/15548627.2015.1063871.

31. Helms, E.J., Berry, M.W., Chaw, R.C., DuFort, C.C., Sun, D., Onate, M.K., Oon, C., Bhattacharyya, S., Sanford-Crane, H., Horton, W., et al. (2022). Mesenchymal Lineage Heterogeneity Underlies Nonredundant Functions of Pancreatic Cancer-Associated Fibroblasts. Cancer Discov 12, 484–501. 10.1158/2159-8290.CD-21-0601.

32. Manoukian, P., Bijlsma, M., and van Laarhoven, H. (2021). The Cellular Origins of Cancer-Associated Fibroblasts and Their Opposing Contributions to Pancreatic Cancer Growth. Front Cell Dev Biol 9, 743907. 10.3389/fcell.2021.743907.

33. Hwang, R.F., Moore, T., Arumugam, T., Ramachandran, V., Amos, K.D., Rivera, A., Ji, B., Evans, D.B., and Logsdon, C.D. (2008). Cancer-associated stromal fibroblasts promote pancreatic tumor progression. Cancer Res 68, 918–926. 10.1158/0008-5472.CAN-07-5714.

34. Elyada, E., Bolisetty, M., Laise, P., Flynn, W.F., Courtois, E.T., Burkhart, R.A., Teinor, J.A., Belleau, P., Biffi, G., Lucito, M.S., et al. (2019). Cross-Species Single-Cell Analysis of Pancreatic Ductal Adenocarcinoma Reveals Antigen-Presenting Cancer-Associated Fibroblasts. Cancer Discov 9, 1102–1123. 10.1158/2159-8290.CD-19-0094.

35. Gupta, S., Yano, J., Mercier, V., Htwe, H.H., Shin, H.R., Rademaker, G., Cakir, Z., Ituarte, T., Wen, K.W., Kim, G.E., et al. (2021). Lysosomal retargeting of Myoferlin mitigates membrane stress to enable pancreatic cancer growth. Nat Cell Biol 23, 232–242. 10.1038/s41556-021-00644-7.

36. Maghe, C., Trillet, K., Andre-Gregoire, G., Kerherve, M., Merlet, L., Jacobs, K.A., Schauer, K., Bidere, N., and Gavard, J. (2024). The paracaspase MALT1 controls cholesterol homeostasis in glioblastoma stem-like cells through lysosome proteome shaping. Cell Rep 43, 113631. 10.1016/j.celrep.2023.113631.

37. Ozdemir, B.C., Pentcheva-Hoang, T., Carstens, J.L., Zheng, X., Wu, C.C., Simpson, T.R., Laklai, H., Sugimoto, H., Kahlert, C., Novitskiy, S.V., et al. (2014). Depletion of carcinoma-associated fibroblasts and fibrosis induces immunosuppression and accelerates pancreas cancer with reduced survival. Cancer Cell 25, 719–734. 10.1016/j.ccr.2014.04.005.

38. Ebner, M., Puchkov, D., Lopez-Ortega, O., Muthukottiappan, P., Su, Y., Schmied, C., Zillmann, S., Nikonenko, I., Koddebusch, J., Dornan, G.L., et al. (2023). Nutrient-regulated control of lysosome function by signaling lipid conversion. Cell 186, 5328–5346 e5326. 10.1016/j.cell.2023.09.027.

39. Eriksson, I., and Ollinger, K. (2024). Lysosomes in Cancer-At the Crossroad of Good and Evil. Cells 13. 10.3390/cells13050459.

40. Wang, F., Gomez-Sintes, R., and Boya, P. (2018). Lysosomal membrane permeabilization and cell death. Traffic 19, 918–931. 10.1111/tra.12613.

41. Zoncu, R., and Perera, R.M. (2022). Built to last: lysosome remodeling and repair in health and disease. Trends Cell Biol 32, 597–610. 10.1016/j.tcb.2021.12.009.

42. Aits, S., and Jaattela, M. (2013). Lysosomal cell death at a glance. J Cell Sci 126, 1905–1912. 10.1242/jcs.091181.

43. Sousa, C.M., Biancur, D.E., Wang, X., Halbrook, C.J., Sherman, M.H., Zhang, L., Kremer, D., Hwang, R.F., Witkiewicz, A.K., Ying, H., et al. (2016). Pancreatic stellate cells support tumour metabolism through autophagic alanine secretion. Nature 536, 479–483. 10.1038/nature19084.

44. van de Vlekkert, D., Demmers, J., Nguyen, X.X., Campos, Y., Machado, E., Annunziata, I., Hu, H., Gomero, E., Qiu, X., Bongiovanni, A., et al. (2019). Excessive exosome release is the pathogenic pathway linking a lysosomal deficiency to generalized fibrosis. Sci Adv 5, eaav3270. 10.1126/sciadv.aav3270.

45. Richards, K.E., Zeleniak, A.E., Fishel, M.L., Wu, J., Littlepage, L.E., and Hill, R. (2017). Cancer-associated fibroblast exosomes regulate survival and proliferation of pancreatic cancer cells. Oncogene 36, 1770–1778. 10.1038/onc.2016.353.

46. Roberts, E.W., Deonarine, A., Jones, J.O., Denton, A.E., Feig, C., Lyons, S.K., Espeli, M., Kraman, M., McKenna, B., Wells, R.J., et al. (2013). Depletion of stromal cells expressing fibroblast activation protein-alpha from skeletal muscle and bone marrow results in cachexia and anemia. J Exp Med 210, 1137–1151. 10.1084/jem.20122344.

47. Chandler, C., Liu, T., Buckanovich, R., and Coffman, L.G. (2019). The double edge sword of fibrosis in cancer. Transl Res 209, 55–67. 10.1016/j.trsl.2019.02.006.

48. Potdar, S., Ianevski, F., Ianevski, A., Tanoli, Z., Wennerberg, K., Seashore-Ludlow, B., Kallioniemi, O., Ostling, P., Aittokallio, T., and Saarela, J. (2023). Breeze 2.0: an interactive web-tool for visual analysis and comparison of drug response data. Nucleic Acids Res 51, W57–W61. 10.1093/nar/gkad390.

49. Stoeckius, M., Zheng, S., Houck-Loomis, B., Hao, S., Yeung, B.Z., Mauck, W.M., 3rd, Smibert, P., and Satija, R. (2018). Cell Hashing with barcoded antibodies enables multiplexing and doublet detection for single cell genomics. Genome Biol 19, 224. 10.1186/s13059-018-1603-1.

