## Supplemental Figures S1-7, TAble S1 for "Therapeutic Eradication of Cancer-associated Fibroblasts Inhibits in vivo progression of Pancreatic Cancer"

^Equal contribution

on

[1]Department of Surgery, Helsinki University Hospital, Helsinki, [2]Translational Cancer Medicine Research Program, Faculty of Medicine, University of Helsinki, Helsinki, [3]iCAN Digital Precision Cancer Medicine Flagship, University of Helsinki, Helsinki, [4]Department Of Surgical Oncology, Division Of Surgery, The University Of Texas MD Anderson Cancer Center, Houston, TX, USA, [5]Neuroscience Center, Helsinki Institute of Life Science (HiLIFE), University of Helsinki, Helsinki, Finland, [6]Laboratory Animal Center, Helsinki Institute of Life Science (HiLIFE), University of Helsinki, Helsinki, Finland.


**Running title:** Clemastine inhibits PDAC progression

### Supplementary Figures

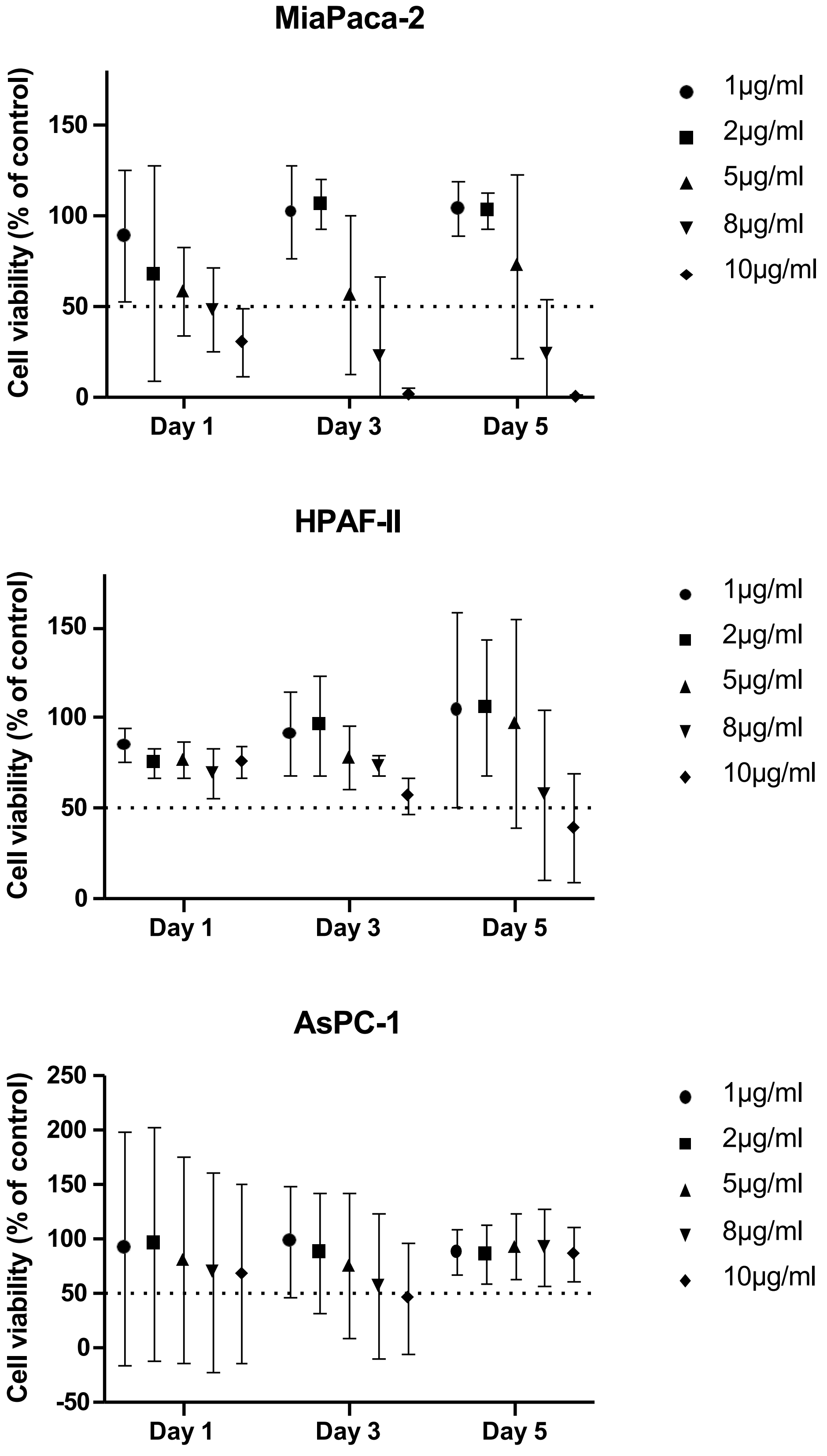


**Supplementary Figure S1.**

**Supplementary Figure S1. Sensitivity of PDAC cell lines to clemastine.** Clemastine treatment of three commercial cell lines (MiaPaca-2, HPAF-II and AsPC-1). Cell viability (mean with 95% confidence intervals) was measured using the XTT assay at the indicated clemastine concentrations and time points. Dashed line marks the 50% cell viability. MiaPaca-2 was the only commercial cell line showing significant sensitivity for clemastine.

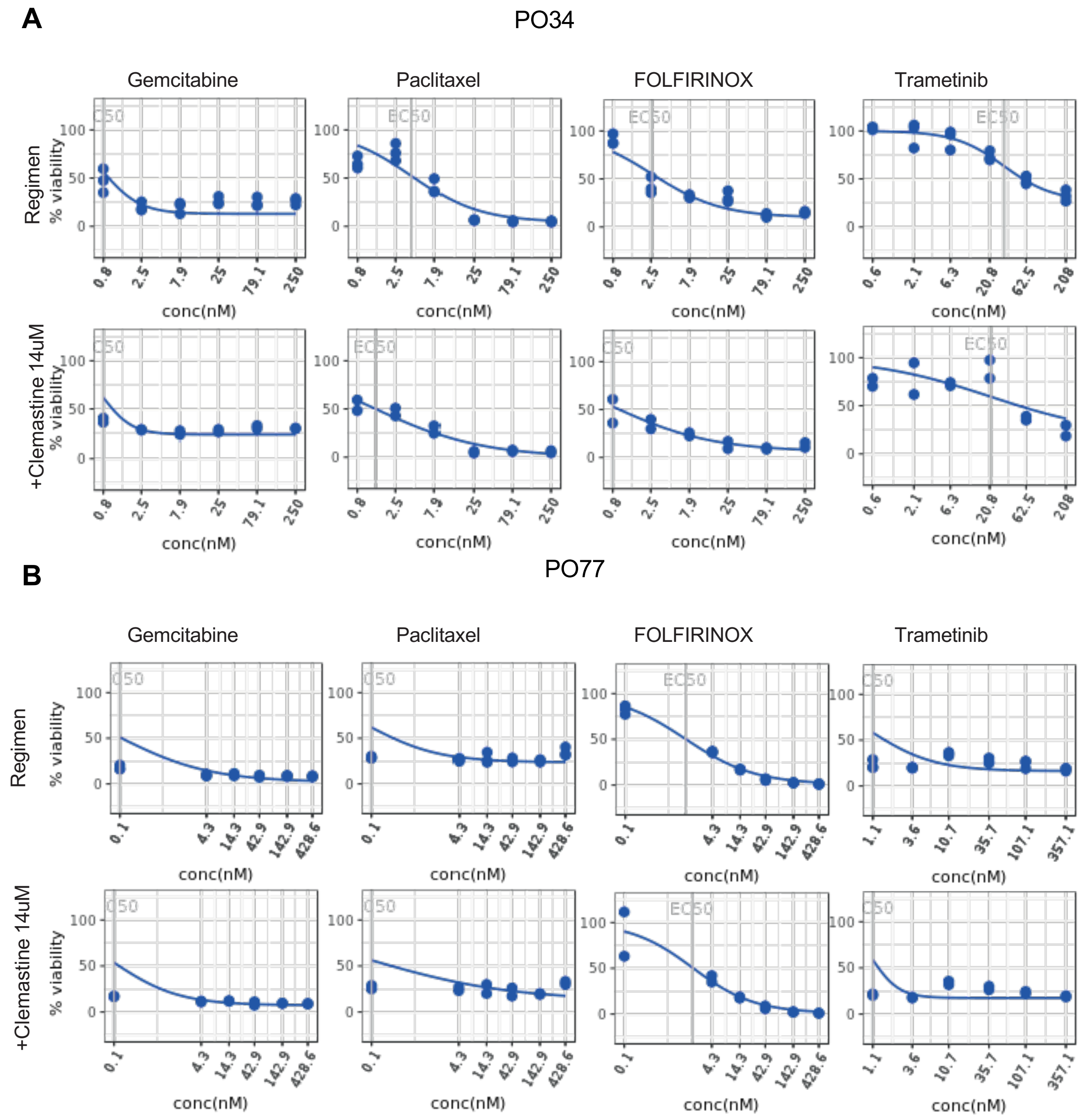


C

| PO | Regimen | +Clemastine | EC 50 | | AU C | | DSS | |
|---|---|---|---|---|---|---|---|---|
| PO34 | Gemcit | abine | 0.8 | 0.8 | 202.6 | 180.7 | 40.8 | 36.8 |
| | Paclit | axel | 4 | 1.4 | 164.6 | 191.5 | 31.4 | 37.1 |
| | FOLFIR | INOX | 2.6 | 0.8 | 167.9 | 193.1 | 32.6 | 38 |
| | Trame | tinib | 31.9 | 21.6 | 61.2 | 87.9 | 10.3 | 14.3 |
| PO77 | Gemcit | abine | 0.1 | 0.1 | 298.9 | 297.7 | 40.3 | 40.6 |
| | Paclit | axel | 0.1 | 0.1 | 243 | 245.3 | 33.6 | 32.9 |
| | FOLFIR | INOX | 1.3 | 1.9 | 239.2 | 228.1 | 31.1 | 29.4 |
| | Trame | tinib | 1.1 | 1.1 | 186.9 | 197.3 | 37.2 | 39.7 |

**Supplementary Figure S2**

**Supplementary Figure S2. Clemastine did not show synergistic effect with four tested regimens in vitro.** To study the possible synergistic effect of clemastine with four PDAC regimens (Trametinib, Paclitaxel, Gemcitabine, and FOLFIRINOX), PDAC organoids (PO) from two patients (PO34 and PO77) were treated with different drug concentrations either alone or in combination with clemastine. Drug response curves for PO34 (**A**) and PO77 (**B**) are shown. **C.** EC50, AUC, and DSS for each regimen alone and in combination with clemastine for both PO34 and PO77. We detected no significant synergy of clemastine with any of the tested drugs.

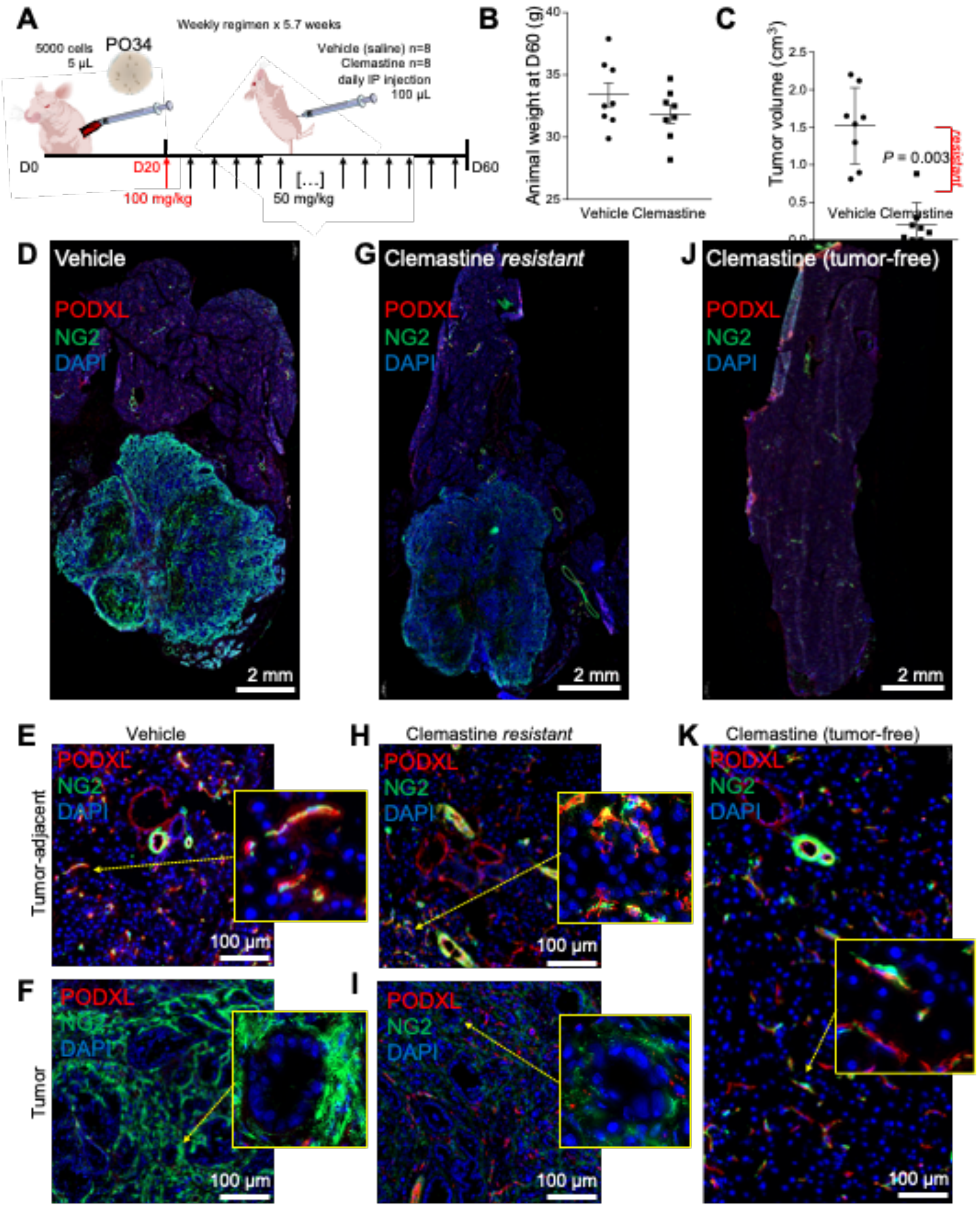


**Supplementary Figure S3**

**Supplementary Figure S3. Preclinical evaluation of clemastine in PO34 PDAC patient avatar model. A**. Summary and timeline of the protocol; orthotopic implantation of patient PDAC cells via the splenic vein, daily intraperitoneal dosage of saline (Vehicle) or clemastine (first day 100 then 50 mg/kg) starting at 20 days post-implantation for 40 days. **B.** Mice body mass at the day of termination of the experiment (D60). **C.** Pancreatic tumour volume ($cm^3$) in the vehicle and clemastine-treated groups. *P*-value calculated with paired *t*-test. **D-K**. Immunofluorescent histological sections of pancreas from the vehicle (D-F), clemastine resistant "outlier" (G-I), and clemastine sensitive (J, K) PO34 patient avatars. Visualization of blood vessels (PODXL, red), cancer-associated fibroblasts (NG2, green), and cell nuclei using DAPI (blue). Non-tumor pancreatic tissue (E, H, K) shows distinct stromal arrangement of blood vessels and fibroblasts compared to the neoplastic areas (F, I).

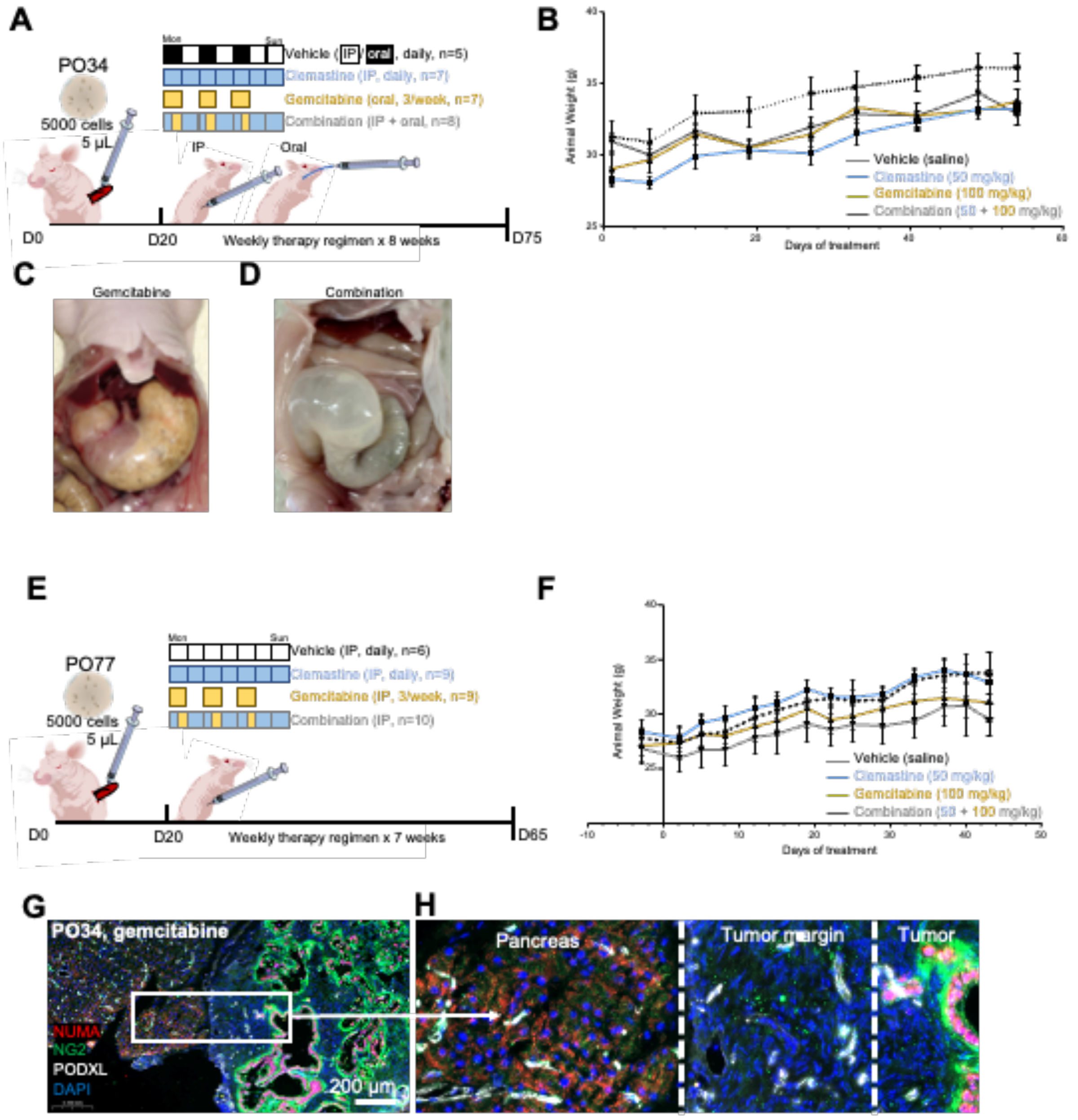


**Supplementary Figure S4**

**Supplementary Figure S4. Preclinical evaluation of clemastine, gemcitabine, and combination treatment in PDAC patient avatar models. A**. Summary and timeline of the protocol; orthotopic implantation of patient PO34 PDAC cells, intraperitoneal dosage of saline (Vehicle, N=5), clemastine (blue, 50 mg/kg, N=7), oral gemcitabine (yellow, 100 mg/kg, N=7) or clemastine+gemcitabine (N=8), starting 20 days post-implantation for 8 weeks. **B**. PO34 patient avatar body mass (g) follow-up in the four treatment groups. **C.** Side-effect of oral gemcitabine (dyspepsia). **D.** Side-effect of the combined dosage (intestinal/cecal inflammation). **E.** Summary and timeline of the protocol; orthotopic implantation of patient PO77 PDAC cells, intraperitoneal dosage of saline (Vehicle, N=6), clemastine (blue, 50 mg/kg, N=9), gemcitabine (yellow, 100 mg/kg, N=9), or clemastine+gemcitabine combinstion (N=10), starting 20 days post-implantation for 8 weeks. **F.** PO77 patient avatar body mass (g) follow-up in the four treatment groups. **G**. Immunofluorescent histological section of pancreas from PO34 patient avatar treated with gemcitabine and stained to visualize human nuclear antigen (NUMA, red), blood vessels (PODXL, white), cancer-associated fibroblasts (NG2, green), and cell nuclei using DAPI (blue). **H.** Higher magnification from the boxed area in panel G, highlighting the distribution of blood vessels between the healthy pancreatic tissue, tumor margins, and NUMA$^{+}$ tumor residuals.

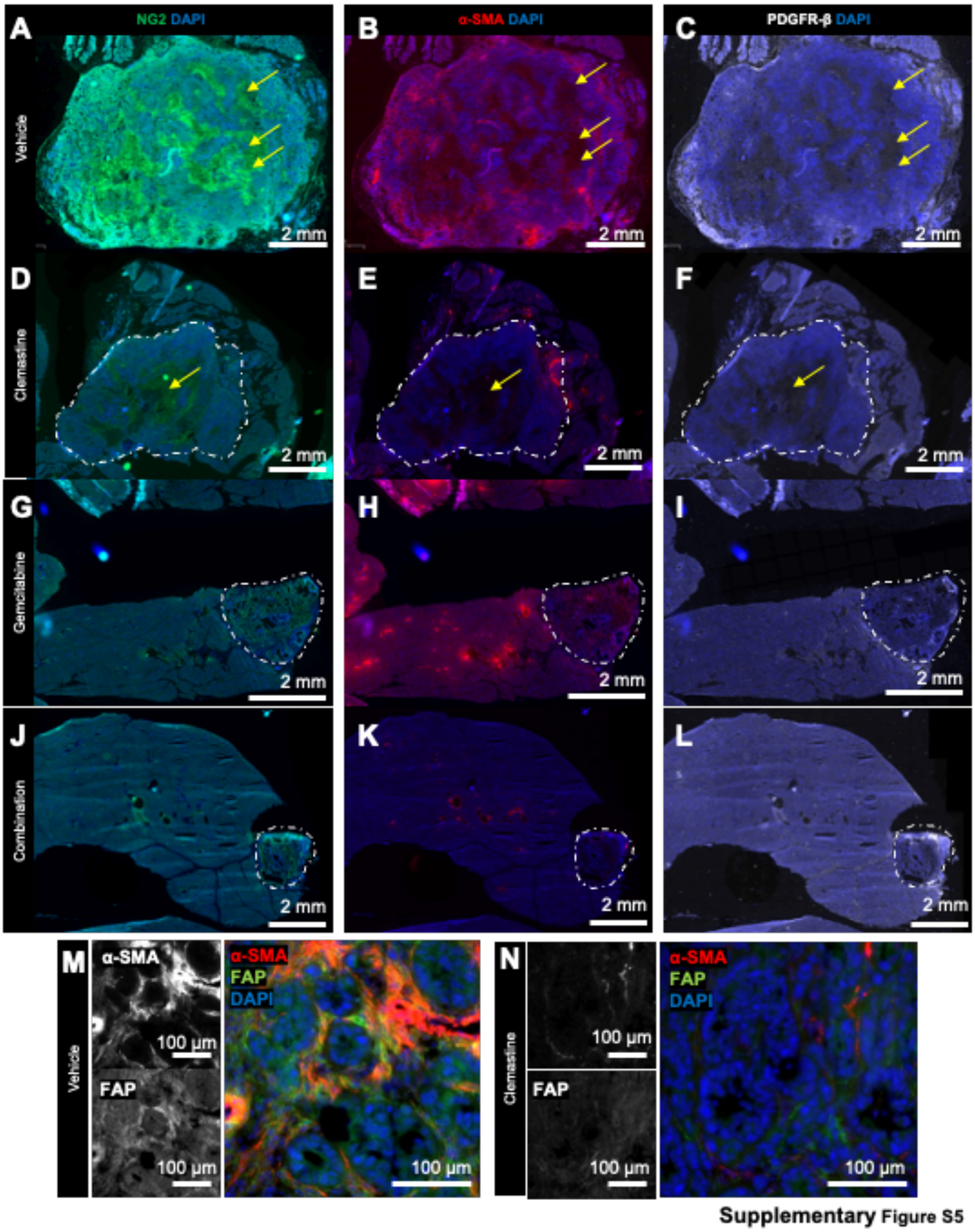


**Supplementary Figure S5. Distribution of cancer-associated fibroblast subtypes in the PO34-derived PDAC patient avatars. A-C.** Whole tissue immunofluorescence micrographs of NG2+ periglandular inflammatory CAFs (A, green), periglandular α-SMA+ (B, red), and pro-metastatic PDGFRβ+ (C, white) myofibroblastic CAFs in vehicle-treated PO34 PDAC patient avatar. **D-F**. Whole tissue immunofluorescence micrographs of NG2+ periglandular inflammatory CAFs (A, green), periglandular αSMA+ (B, red), and pro-metastatic PDGFRβ+ (C, white) myofibroblastic CAFs in clemastine-treated PO34 PDAC patient avatar. **G-I.** Whole tissue immunofluorescence micrographs of NG2+ periglandular inflammatory CAFs (A, green), periglandular αSMA+ (B, red), and pro-metastatic PDGFR β+ (C, white) myofibroblastic CAFs in gemcitabine-treated PO34 PDAC patient avatar. **J-L**. Whole tissue immunofluorescence micrographs of NG2+ periglandular inflammatory CAFs (A, green), periglandular αSMA+ (B, red), and pro-metastatic PDGFRβ+ (C, white) myofibroblastic CAFs in clemastine/gemcitabine combination-treated PO34 PDAC patient avatar. Dotted lines delineate the tumor in the pancreas, yellow arrows highlight necrotic areas, cell nuclei were counterstained with DAPI (blue). **M**. Immunofluorescence micrographs of periglandular αSMA+ (red) and FAP+ inflammatory CAFs (green) in vehicle-treated PO34 tumors. **N**. Immunofluorescence micrographs of periglandular αSMA+ (red) and FAP+ inflammatory CAFs (green) in clemastine-treated PO34 tumors.

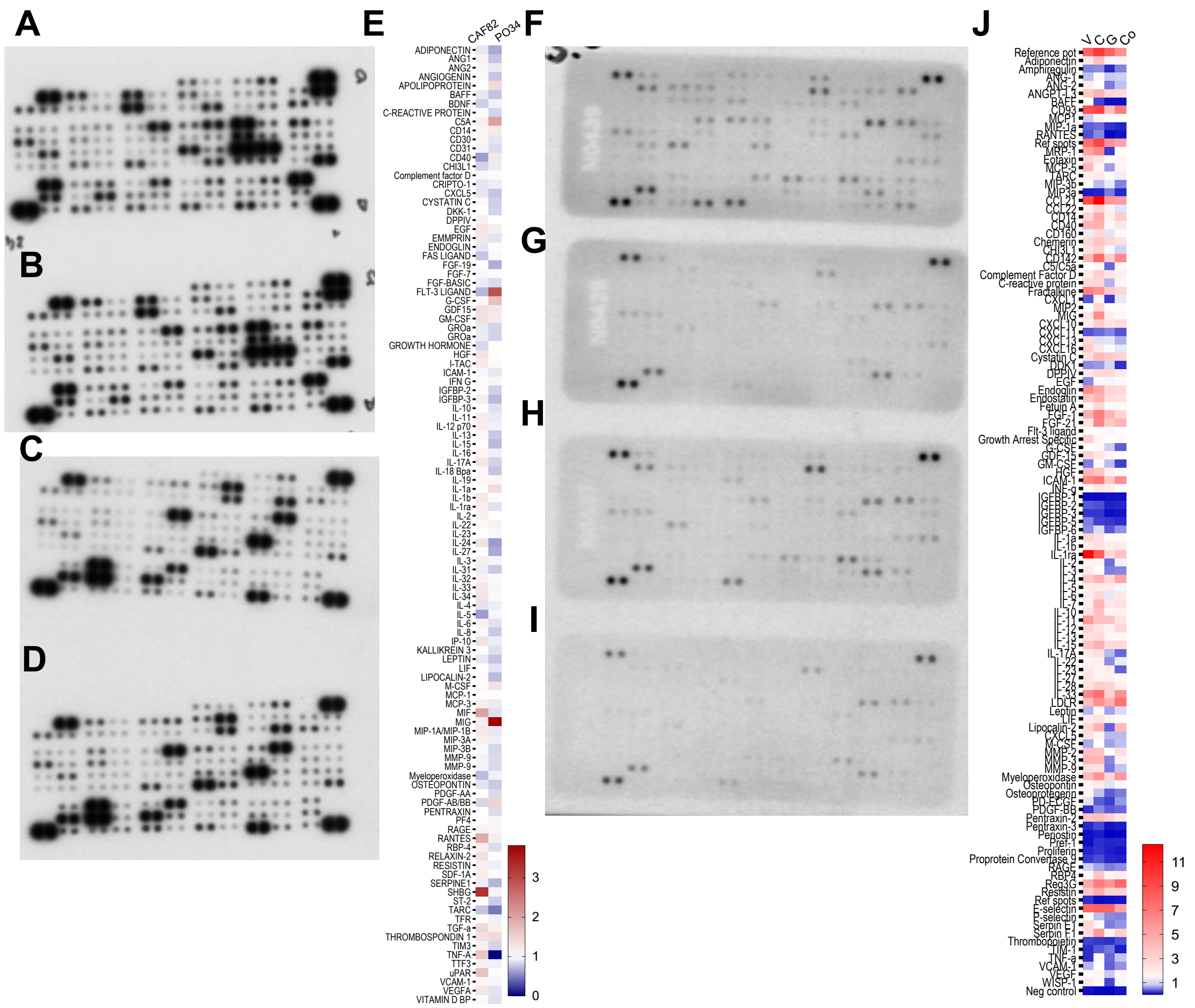


**Supplementary Figure S6**

**Supplementary Figure S6. Raw data for the dot blot secretome arrays. A.** Conditioned medium from human PDAC derived CAF82 cells was analyzed using the human cytokine array kit to study the secreted proteins in a chemiluminescent dot blot. **B.** Conditioned medium from clemastine-treated (2 ug/mL, 18 h) human PDAC derived CAF82 cells analyzed using the human cytokine array kit. **C**. Conditioned medium from human PDAC organoid PO34 cells analyzed using the human cytokine array kit. **D.** Conditioned medium from clemastine-treated (2 ug/mL, 18 h) human PDAC organoid PO34 cells analyzed using the human cytokine array kit. **E**. Quantification of the dot blots. Data are shown as mean of fold-over control untreated cells. **F**. Whole tissue extracts from the pancreas of PO77 patient avatar treated with vehicle were analyzed using the mouse cytokine array kit to study the secreted proteins in a chemiluminescent dot blot. **G.** Whole tissue extracts from the pancreas of PO77 patient avatar treated with clemastine were analyzed using the mouse cytokine array kit. **H.** Whole tissue extracts from the pancreas of PO77 patient avatar treated with gemcitabine were analyzed using the mouse cytokine array kit. **I.** Whole tissue extracts from the pancreas of PO77 patient avatar treated with combination of clemastine and gemcitabine were analyzed using the mouse cytokine array kit. **J.** Quantification of the dot blots. Heatmap of PO77 PDAC patient avatars from the vehicle (V), clemastine (C), gemcitabine (G), and combination (Co) treatment groups. Data are shown as mean protein dot intensity value (a.u.) normalized to the assay reference dots.

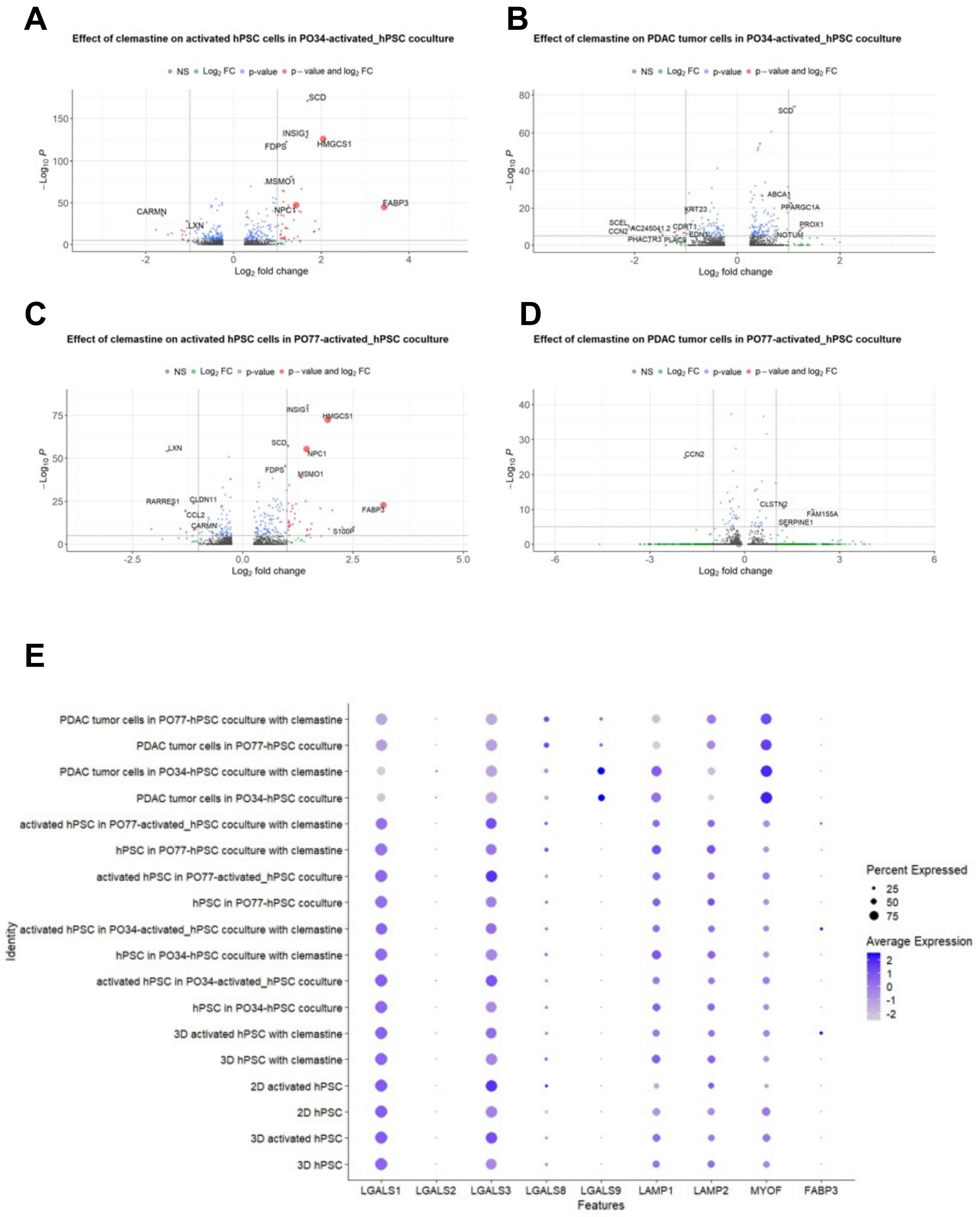


**Supplementary Figure S7**

**Supplementary Figure S7. Single-cell transcriptomic profiling of PDAC organoids and hPSC co-cultures. A-B.** Volcano plots showing down- and upregulated genes in the clemastine treated co-cultures of PO34 and hPSCs. **C-D**. Volcano plots showing down- and upregulated genes in the clemastine treated co-cultures of PO77 and hPSCs. **E**. Dot plot shows the expression of LGALS1, 2, 3, 8, and 9 as well as LAMP1, 2, MYOF, and FABP3 in tumor cells and in the pancreatic stellate cells (fibroblasts).

**Supplementary Table 1.** Concentrations of clemastine, trametinib, paclitaxel, gemcitabine and FOLFIRINOX (nM) and the four compounds and their relative concentrations in FOLFIRINOX.

| | Clemastine (nM) | Trametinib (nM) | Paclitaxel (nM) | Gemcitabine (nM) | FOLFIRINOX |
|---|---|---|---|---|---|
| 1 | 2907.9 | 208 | 250 | 250 | 3000 |
| 2 | 5815.8 | 62.5 | 79.1 | 79.1 | 950 |
| 3 | 11631.6 | 20.8 | 25 | 25 | 300 |
| 4 | 17447.4 | 6.3 | 7.9 | 7.9 | 95 |
| 5 | 23263.3 | 2.1 | 2.5 | 2.5 | 30 |
| 6 | 29079.1 | 0.6 | 0.8 | 0.8 | 9.5 |

| FOLFIRINOX | Concentration (nM) |
|---|---|
| 5-FU | 3009.4 |
| SN-38 | 451.4 |
| Leucovorin | 812.5 |
| Oxaliplatin | 209.4 |
